# Flood-pulse disturbances as a threat for long-living Amazonian trees

**DOI:** 10.1101/2019.12.18.872598

**Authors:** Angélica F Resende, Maria T F Piedade, Yuri O Feitosa, Victor Hugo F Andrade, Susan E Trumbore, Flávia M Durgante, Maíra O Macedo, Jochen Schöngart

## Abstract

The long-living tree species *Eschweilera tenuifolia* (O. Berg) Miers (Lecythidaceae) is characteristic to oligotrophic floodplain forests (*igapó*) influenced by a regular and predictable flood-pulse. This species preferentially occurs at macrohabitats flooded up to 10 months per year forming monodominant stands. We aimed to analyze the growth and mortality patterns of this species under pristine conditions (Jaú National Park-JNP) and in an impacted *igapó* (Uatumã Sustainable Development Reserve-USDR) where the downstream flood-pulse disturbance occasioned by the Balbina hydroelectric plant caused massive mortality of this species. Using a total of 91 trees (62 living and 29 dead) at the USDR and 52 (31 living and 21 dead) from JNP, we analyzed age-diameter relationships, mean passage time through 5-cm diameter classes, growth change patterns, growth ratios, clustering of mean diameter increment (MDI), and dated the year of death from each individual using radiocarbon (^14^C) analysis. Growth and mortality patterns were then related to climatic or anthropogenic disturbances. Our results show similar structural parameters for both studied populations regarding the estimated maximum ages of 466 years (JNP) and 498 years (USDR) and MDI, except for one single tree at the USDR with an estimated age of 820 yrs. Living trees from JNP showed distinctly altered growth after 1975, probably related to consecutive years of high annual minimum water levels. Tree mortality in the JNP occurred during different periods, probably induced by extreme hydroclimatic events. At the USDR changes in growth and mortality patterns occurred after 1983, when the Balbina dam construction started. Despite being one of the best flood-adapted tree species, *E. tenuifolia* seems to be sensitive to both, long-lasting dry and wet periods induced by climatic or anthropogenic disturbances or resulting synergies among both. Even more than 30 years after the start of disturbances at the USDR, the flood-pulse alteration continues affecting both mortality and growth of this species which can potentially cause regional extinction.

## Introduction

In recent decades significative disturbances of the flood-pulse regimes have been observed as a consequence of climate change leading to an increase in the frequency and magnitude of extreme floods in Central Amazonia (Barichivich et al., 2018) and due to unprecedented land-use changes, such as the construction of many hydroelectric power plants (Latrubesse et al., 2017; Lees, Peres, Fearnside, Schneider, & Zuanon, 2016) strongly affecting the entire hydrological regimes (Assahira et al., 2017; Timpe & Kaplan, 2017). The hydrological cycle of the large Central Amazonian rivers is characterized by monomodal flood-pulses of high amplitudes with a high and low water period during the year inducing a distinct seasonality in the associated floodplains covering over 750.000 km^2^ (Wolfgang J. Junk, Bayley, & Sparks, 1989; Wolfgang J. Junk et al., 2011; Wittmann & Junk, 2016). The Amazon basin hosts one of the last remaining networks of free-flowing rivers, among them four of the ten largest rivers on Earth, representing about 18% of the global freshwater discharge to the oceans (Grill et al., 2019; Latrubesse, 2008).

Black-water rivers (e.g., Negro and Uatumã rivers) draining the Precambrian Guyana Shield form acidic, sediment-poor Amazonian floodplains with a high forest cover (>85%) are called *igapó* and comprise over 140,000 km^2^ of the Amazon basin (Wolfgang J. Junk, Wittmann, Schöngart, & Piedade, 2015; Melack & Hess, 2010). These environments are older and geomorphologically more stable compared to the dynamic *várzea* along the sediment-loaded white-water rivers originating from the Andes (Furch, 1997; Prance, 1980; Sioli, 1984). Both floodplain types are subject to annual flood pulses (Wolfgang J. Junk et al., 1989), causing inundations which can last from a couple of weeks to more than eight months in the várzea and even up to ten months in the igapó (Wittmann, Schöngart, & Junk, 2010), depending on the topography of each floodplain macrohabitat.

Floodplain tree species feature sophisticated anatomical, morphological, physiological and biochemical adaptations that allow them to overcome the anaerobic conditions induced by flooding (Parolin, 2009; Parolin et al., 2004; M. T. F. F. Piedade et al., 2010). A set of different adaptations can be found usually acting in combination, among them the production of phytohormones such as ethylene, directly linked to root elongation, formation of aerenchyma, adventitious roots and hypertrophied lenticels as well as rhizodermal incorporation of suberin, anaerobic pathways of metabolisms and elimination of phytotoxic compounds (Armstrong & Drew, 2002; Crawford & Braendle, 2007; Jackson, 1990; Parolin, 2009; Sauter, 2013; Voesenek, 2003). These adaptations have been developed due to the evolutionary pressure triggered by the flood-pulse over millions of years (W J Junk, 1989; Wittmann et al., 2010) resulting in the most tree species diverse floodplain forests worldwide with a high percentage of endemics (Wittmann *et al*. 2006, 2013).

The anoxic conditions induced by the annual flood-pulse lead to a decrease in diameter increment and a cambial dormancy during the high-water period resulting in the formation of annual tree rings (Schöngart, Piedade, Ludwigshausen, Horna, & Worbes, 2002; Worbes, 1989) and ring width shows a positive correlations with the duration of the non-flooded period (Batista & Schöngart, 2018; Schöngart et al., 2004; Schöngart, Piedade, Wittmann, Junk, & Worbes, 2005). However, *igapó* species grow slower than *várzea* species because of the oligotrophic condition of soil and water (Da Fonseca Júnior, Piedade, & Schöngart, 2009; Rosa et al., 2017; Schöngart et al., 2005; Worbes, 1997). Tree-rings studies reveal by retrospective analyses information on the species’ ecology and allows to detect disturbances in the past, caused by exogenous (e.g., fires, severe droughts, floods) or endogenous (inter-tree competition) events (Assahira et al., 2017; Baker, Bunyavejchewin, Oliver, & Ashton, 2005; Cook & Kairiukstis, 1990; Schweingruber, 1996; Speer, 2009).

In the Amazon region, severe droughts and floods are not a recent phenomenon, since they have also been registered in the past (Granato-Souza et al., 2019; Marengo et al., 2016). During the last three recent decades, however, an intensification of extreme hydroclimatic events has been observed, especially of extreme flood events in the Central Amazon region (Barichivich et al., 2018; Gloor et al., 2013), while in the southern part of the Amazon region an increasing length of the dry season is evident (Marengo et al., 2018). The causes for hydroclimatic extreme events (droughts and floods) are mainly the *El Niño*-Southern Oscillation (ENSO) originating from the Equatorial Pacific and the Meridional Mode across the tropical Atlantic Ocean determining the position of the Intertropical Convergence Zone (ITCZ) (Nobre, Marengo, & Artaxo, 2009), both leading to extreme hydroclimatic events at different spatial and temporal scales across the Amazon basin (Marengo and Espinoza 2016). Since climate and hydrology are the foremost drivers of wetlands formation, the recent intensification of both factors is probably affecting the floodplain forests, especially the most flood-adapted tree species. Furthermore, massive land-use changes, such as the expansion of hydroelectric power plants, mining, urban footprints, deforestation for livestock and agricultural production are growing vectors of disturbances in the Amazon basin. Synergies of climate and land-use changes are growing threats for its megadiversity and ecosystem functioning, especially for the hydrological cycle which is intrinsically related to the carbon balance (Artaxo, 2019; Castello et al., 2013; Davidson et al., 2012; Fearnside, 1990b, 2005; Lovejoy & Nobre, 2018; Malhi et al., 2008; Nobre et al., 2016). Concerning hydroelectric power plants, the flood-pulse loss by damming rivers affects the riparian ecosystems downstream of the dam (Ligon, Dietrich, & Trush, 1995; Lobo, Wittmann, & Piedade, 2019; Neves, Piedade, Resende, Feitosa, & Schöngart, 2019), causes massive tree mortality (Assahira et al., 2017; Angélica Faria de Resende et al., 2019) and affects the aquatic fauna (Lees et al., 2016).

Monodominant and opened stands at the most inundated *igapó* environments are colonized by the tree species *Eschweilera tenuifolia* (Lecythidaceae) (Wolfgang J. Junk, Piedade, Cunha, Wittmann, & Schöngart, 2018; Wolfgang J. Junk et al., 2015; S. A. Mori, 2001; ter Steege et al., 2019). The harsh environmental conditions and low hydrological buffer capacity turn this macrohabitat vulnerable for these populations against substantial changes in the flood-pulse regime caused by both, anthropogenic and hydroclimatic, disturbances (Wolfgang J. Junk et al., 2018, 2015; Angélica Faria de Resende et al., 2019). In this study, we analyze growth and mortality patterns of two *E. tenuifolia* populations using dendroecological tools and radiocarbon dating to elucidate the impact of hydrological disturbances associated with changes in the magnitude and frequency of hydrological alterations affecting the species’ growth and survival. We compare a population growing under pristine conditions in the *igapó* along the Jaú River (Jaú National Park - JNP) to another one growing in the *igapó* of the Uatumã Sustainable Development Reserve (USDR), downstream of the hydroelectric Balbina dam which was established in the 1980s damming the Uatumã River, causing massive disturbances in the hydrological regime (Assahira et al., 2017). In this study, we test if diameter growth patterns differ between and within both systems during pristine conditions and after the alteration of the hydrological cycle caused by the dam implementation (USDR) and the period of flood-pulse intensification (JNP. Furthermore, we date the year of death of trees in both regions using radiocarbon dating to link mortality patterns to natural and anthropogenic disturbances.

## Methods

### The tree species Eschweilera tenuifolia (O. Berg) Miers

*Eschweilera tenuifolia* belongs to the Amazonia nut family Lecythidaceae and is commonly known as *macacarecuia* (*macaco* means monkey and *cuia* means gourd, in Brazilian Portuguese) or *cuieira* (what means gourd tree). The genus *Eschweilera* is the biggest among the Lecythidaceae family, occurring in the most different environments. The species *E. tenuifolia* is the only one in the section *Jugastrum* (*E. parvifolia* clade) (Huang, Mori, & Kelly, 2015) because even having a flower typical of *Eschweilera* it lacks a flower pedicel, has unique wedge-shaped seeds, the aril is absent, there is a corky seed coat, and seed germination is lateral. The fruit is bowl-shaped (*cuia*), and mature trees are about 15 to 20 meters tall with diameters of up to two meters (Huang et al., 2015; S. A. Mori & Prance, 1990).

Despite the hard-shelled fruits, the seeds are consumed mainly by birds and fishes during the aquatic phase (Maia & Piedade, 2000) and by birds, monkeys, and other terrestrial animals in the terrestrial phase, when is possible to access the trees by land for a short period (Barnett et al., 2002; Barnett, Bowler, Bezerra, & Defler, 2013; Barnett, Castilho, Shapley, & Anicácio, 2005). Leaves and flowers are also consumed when fruits and seeds are scarce (Barnett et al., 2013). The fruits and seeds present sophisticated dispersion adaptations, such as a porous layer and water-repellency that make them buoyant (Kubitzki & Ziburski, 1994). The species belongs to the evergreen ecotype and presents peak leaf abscission during the high water and insolation period, from June to August, while the peak production of new leaves is between August and September (Maia & Piedade, 2000) when the water level is decreasing and the growing period starts for floodplain trees (Schöngart et al. 2002). Flowers and fruits are produced after the new leaves, as it has been described for trees along the Tarumã-mirim River in Central Amazonia (Maia & Piedade, 2000), a Negro River sub-basin where the flooding pattern is similar to our study areas.

*E. tenuifolia* occurs in macrohabitats characterized by mixed forests with high interspecific competition for resources at low topographies in the *igapó* (main river tributaries - Figure S1). However, its main macrohabitat are the lowest level of the floodplain (up to 10 months of flooding), mainly in lakes with clayish substrates, where the species occur in monodominant or even monospecific formation(Lv Ferreira & Prance, 1998; Wolfgang J. Junk et al., 2018, 2015) growing with low inter-tree competition (Schöngart, Bräuning, Barbosa, Lisi, & de Oliveira, 2017) (Figure S2). This species is very abundant throughout the floodplains of the Amazon and Orinoco basins (Lv Ferreira & Prance, 1998; S. A. Mori & Prance, 1990). The wood of the species is dense (0.75 g cm^3^) (Parolin & Ferreira, 1998; Parolin & Worbes, 2000) and larger individuals always present hollows, degraded tree crowns, longitudinally twisted stems and bark-covered knots which is typical for ancient trees (Schöngart, Bräuning, et al., 2017). Therefore *E. tenuifolia* is an interesting bioindicator to be used to analyze past environmental extremes exploring tree-ring analyses.

### Study areas

#### Jaú National Park

The Jaú National Park (JNP) was created in 1980 and it is currently listed as a World Natural Heritage Site and Biosphere Reserve by the United Nations Educational, Scientific and Cultural Organization - UNESCO (Figure 1). It is managed by the Chico Mendes Institute for Biodiversity Conservation - ICMBio. The federal conservation unit comprises 2,272,000 ha situated in the state of Amazonas and has an annual average temperature about 27.4°C and average annual rainfall of about 2215 mm seasonally distributed with maximum 283 mm in April and minimum 96 mm in September (for the period 1983-2018), CRU TS4.03 data from KNMI (Jones & Harris, 2008; Trouet & Van Oldenborgh, 2013). The Jaú River is one of the main tributaries of the Negro River and is characterized by blackish, acidic and nutrient-poor water (L. V. Ferreira, 2000), typical for black-water rivers (Junk et al., 2015). The lower part of its section is mostly affected by the backwater effect of the Negro River. Main vegetation types comprise upland forests (*terra firme*), *igapós*, white-sand forest ecosystems (*campinarana*) and poorly-drained swamps with the predominance of palm trees (*chavascais, buritizais*), forming a mosaic of landscapes due to its varied geological composition and soil drainage (FVA, 1998). The JNP is not considered impacted by anthropic activities, since the few groups of residents that live inside the conservation unit use its resources only for subsistence, without causing significant impacts to the local ecosystem (FVA, 1998).

**Figure 1.**
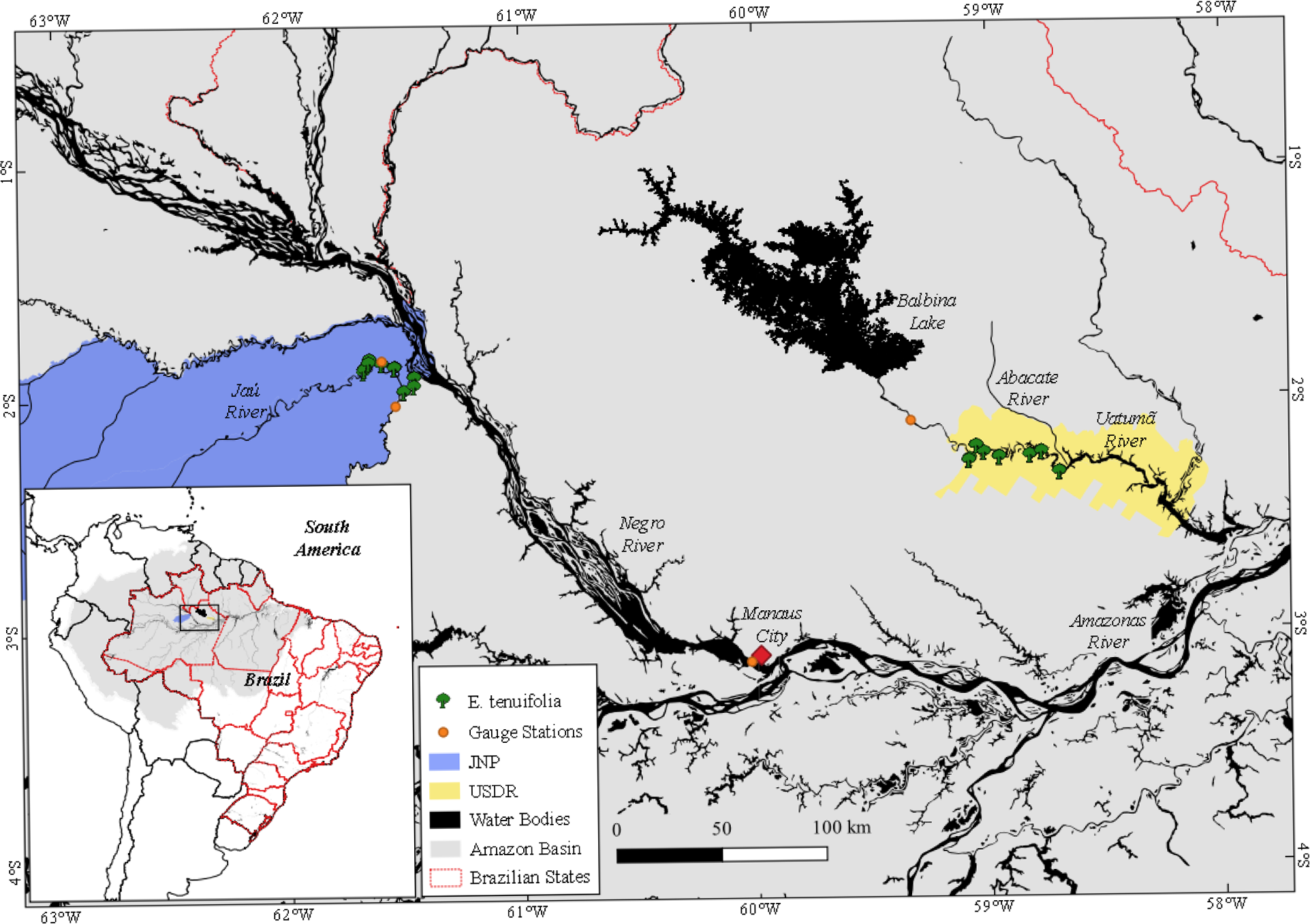
Map showing the location of both study areas in the Jaú National Park (JNP; green) and the Uatumã Sustainable Development Reserve (USDR; pink).

#### Uatumã Sustainable Development Reserve

The Uatumã Sustainable Development Reserve (USDR) was created in 2004 and comprises 424,430 ha located in the State of Amazonas State, about 150 km northeast from the capital Manaus (IDESAM, 2009) (Figure 1). The state conservation unit is composed of different forest ecosystems such as upland forests (*terra firme*), floodplain forests (*igapó*) as well as white- sand ecosystems with opened (*campina*) and closed canopy (*campinarana*) (Andreae et al., 2015). The average annual temperature is around 27.4°C and rainfall is about 2040 mm seasonally distributed with a maximum of 290 mm in March and a minimum of 57 mm in September (for the period 1983-2018, CRU TS4.03 data from KNMI, Jones and Harris, 2008, Trouet and Van Oldenborgh, 2013). The USDR is located downstream of the Balbina Hydroelectric Power Plant, which was installed into the Uatumã River in the period 1983-1987 causing substantial alterations of the hydrological cycle as a result of power generation (Assahira *et al*., 2017). In comparison to the pristine conditions, annual minimum (maximum) water levels increased (decreased) and lost the predictable flood-pulse pattern. The downstream impact in the *igapó* floodplains is severe due to losses of macrohabitats at both extremes due to the suppression of the terrestrial phase (lower elevations) and aquatic phase (upper elevations) (Assahira *et al*., 2019, Lobo *et al*., 2019, Resende *et al*., 2019, Rocha *et al.*, 2019). At the lowest topographies, large-scale mortality of flood-adapted tree species occurs such as *Macrolobium acaciifolium* (Benth.) Benth and *E. tenuifolia* (Assahira et al., 2017; Angélica Faria de Resende et al., 2019). Although the hydrological regimes during pristine conditions (1973-1982) are significantly correlated between the Uatumã and Negro rivers (*R*^2^=0.52; *p*<0.001; Assahira et al. 2017), both have different sizes and geomorphologies of catchments, located in distinct geographic regions of the northern Amazon basin (Figure 1 and Figure S3).

### Data collection in the field

A total of 93 living trees (31 at JNP and 62 at USDR) were sampled in *igapós* along the Jaú and Uatumã rivers, respectively, with diameters at breast height (DBH) varying from 10 to 120 cm (except for one individual with 175 cm at USDR) to consider possible ontogenetic effects on tree growth (Bowman, Brienen, Gloor, Phillips, & Prior, 2013). At least two cores were sampled from each tree using an increment borer of 5 mm internal diameter (except for additional stem discs from two trees of the USDR). From smaller individuals with no hollows, we sampled the entire radius which included the pith, while from larger trees with hollows we collected samples from the intact part of the trunk. Additionally, we sampled cross sections from 21 dead trees in JNP and 29 in USDR using a chainsaw. Further details can be found in the first section of supplementary material.

#### Dendroecological analyses: Diameter growth models and age estimates of hollow trees

Tree rings were measured to calculate the mean annual diameter increment (MDI) per core and individual (average from two or more cores). Average and standard deviation of MDI, DBH, and age for each area were calculated to observe the differences between the same species growing in natural (JNP) and disturbed (USDR) environments. While DBH and MDI were obtained directly from the measured samples, the age had to be estimated for several individuals presenting hollow trunks. Therefore, we produced a preliminary age-DBH relationship by cumulative diameter growth curves based on the ring-width measurements from samples which presented the pith at least on one core. These ring-widths were fitted to the measured DBH in the field by adjusting the measured length of the core to the DBH, proportionally increasing or decreasing the measured ring widths by the ratio obtained dividing the half of the DBH (radius) by the length of the measured core from pith to bark. To account for bark thickness, we obtained this value from cores and cross sections including bark which was then related to the DBH of these individuals. As no significant relationship was observed we used the average bark thickness of 2.35 cm (±1.21 standard deviation) which was subtracted from all measured DBHs of living trees. The constructed and adjusted cumulative diameter growth curves were then fitted to a non-linear (sigmoidal) regression model (Schöngart 2008) presenting well-distributed residuals (Beissner, Genske, Prall, & Weinholz, 2002; R Core Team, 2018):

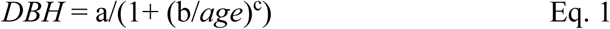

Where DBH is the cumulative diameter for the given year, age the respective age at the given year and *a*, *b*, and *c* are the model parameters.

The diameter of the hollow trunk fraction was estimated, subtracting the average length of the analyzed cores (transformed into diameter) from the measured DBH in the field, corrected by bark thickness (Figure S4). The age of the hollow fraction was then estimated by its diameter using the age-diameter growth model (Eq. 1). The cumulative diameter growth of these individuals started at the estimated age of the hollow trunk diameter. Based on the measured ring-width averaged across the analyzed cores cumulative diameter growth was reconstructed, adjusted for the measured and corrected DBH resulting in an estimated age for these trees. Errors in age estimates caused by irregular shapes of hollows or eccentricity of the pith were reduced by averaging two to four radii for each individual (Figure S4). The procedure was performed for all living and dead trees. Only for the largest tree from USDR with a DBH of 175 cm, we were not able to estimate the age of the large hollow trunk section (diameter of 145 cm) as no other individual from this site presented a DBH larger than 93 cm. To estimate the age of this individual we used a linear regression model based on the MDI of this individual.

#### Median MDI time of diameter growth rates

To analyze growth patterns over the lifespan, we calculated based on the individual cumulative diameter growth curves the MDI through 5-cm diameter classes based on the measured ring-width. Incomplete 5-cm diameter classes originating from hollows or outermost parts of the diameter were not considered. For instance, if a tree of 17.4 cm DBH has a hollow of 2.3 cm in diameter only the size classes 5-10 cm and 10-15 cm were accounted (Figure S5). We then calculated the median and standard deviation of the MDI for each 5-cm size class differentiating dead trees from living trees in both areas (Figure 3) testing the differences through the notched boxplot. Additionally, we performed a Wilcox test (Bauer, 1972) per class comparing macrohabitats within the areas to assure we could combine samples from both macrohabitats (lakes and tributaries). Differences were found only for the classes 15-20 cm at USDR and 10-15 cm at JNP, we considered these differences relatively small, allowing us to combine trees from lakes and tributaries for this analysis.

**Figure 3.**
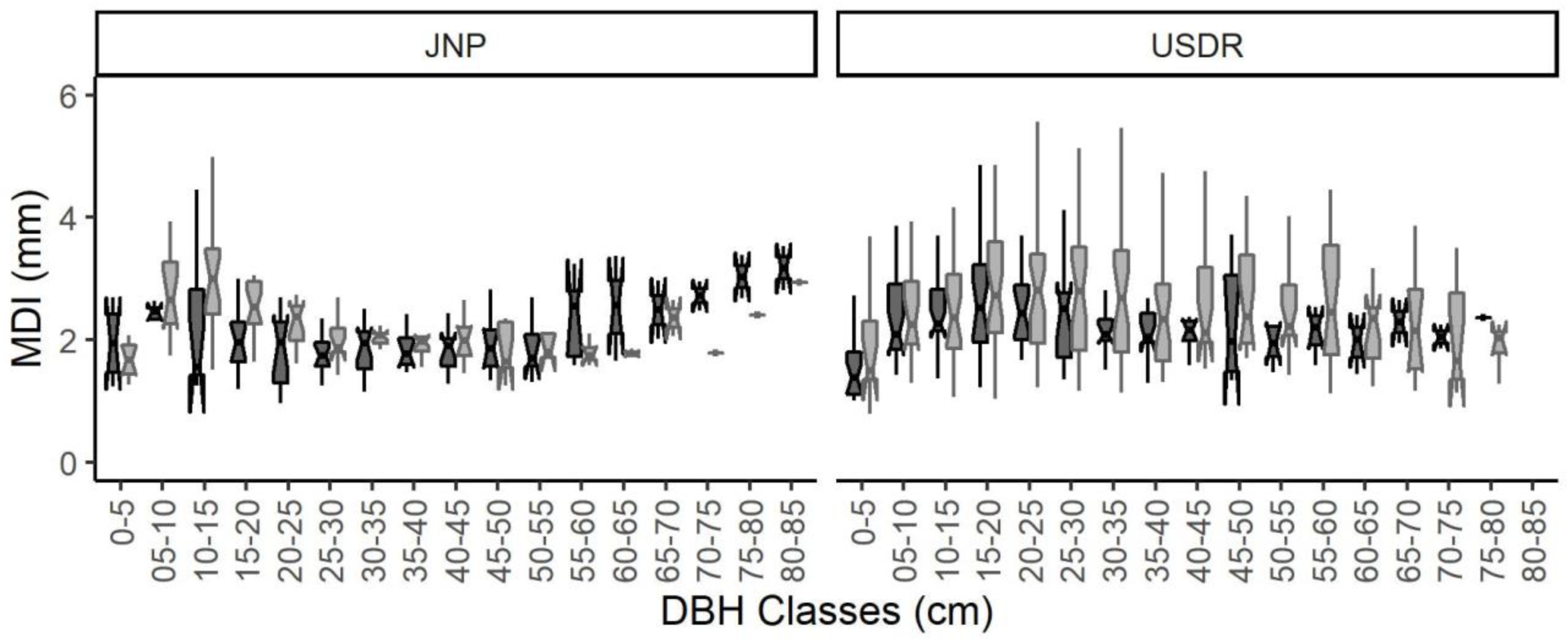
Median MDI across 5-cm diameter (DBH) classes for the Jaú National Park (JNP, left) and the Uatumã Sustainable Development Reserve (USDR, right panel). Living trees are indicated in light grey and the dead ones in black. The notched boxplots are showing the median, 25th and 75th percentiles, as well as the confidence interval (95% confidence) in the notches.

#### Suppression and liberation events and the reduction in growth before death

The raw data of every individual tree-ring series was transformed into relative growth changes *(%GCi*) as suggested by Nowacki and Abrams (1997):

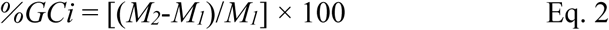

This parameter is obtained for every tree ring (*i*) along the series by the difference between the mean increment (mm/yr) of its subsequent ten years (*M*_*2*_ = *mean increment over years i+1 to i+10*) and the previous ten years (*M*_*1*_ = *mean over years i to i-9*), divided by the mean of the last ten years (*M*_*1*_). By definition, the first and last 10 years of the series are not considered in Eq. 2. This approach has been used to detect the occurrence of suppression and liberation events during canopy attainment in competitive environments such as tropical upland forests (Brienen and Zuidema 2006, Brienen et al. 2010, Schöngart et al. 2015). We used this method to interfere to disturbances in the past affecting tree growth of *E. tenuifolia* trees in both macrohabitats, the open monodominant formation and closed mixed forests, where the principal causes of suppression and release events might be associated to exogenous disturbances (pluriannual severe hydroclimatic events at the monodominant and opened formation on lowest topographies in lakes) or endogenous disturbances resulting from competition for light (closed forests on low topographies along small tributaries). Especially for trees in the USDR, we expected distinct and synchronous changes in tree growth due to the massive disturbance of the hydrological cycle caused by the construction and operation of the hydroelectric dam (Assahira *et al.*, 2017). To define release and suppression events, we established thresholds of *%GC* ≥100% and ≤-50% over at least five consecutive years, respectively (Brienen et al., 2010; Schöngart et al., 2015). The number of trees presenting liberation and suppression events were calculated for each area, considering the tree status (living or dead) and the macrohabitat (lake or tributary).

To understand the “*causa mortis*” of trees, we examined the last formed tree rings of all dead trees to detect patterns of growth declines before mortality occurred based on the method proposed by Cailleret *et al*. (2017). Growth reduction was calculated by the growth ratio (*g*_*r*_) expressed for the diameter increment in a given year of a dead tree in relation to the MDI of living trees in the population with similar DBHs (less than 2 cm of difference) in the same year. A value of *g*_*r*_≤1 is considered a mortality pattern when it occurred immediately before death. The duration of the decay pattern is obtained based on the number of consecutive years before death with *gr* ≤1.

#### Cluster analysis and periodical diameter growth performance

The general dissimilarity coefficient calculated from Gower’s formula (Gower, 1971), was used to estimate the dissimilarity between years using tree-ring widths per study site, by using the R cluster package (Maechler, Rousseeuw, Struyf, Hubert, & Hornik, 2019). The Gower coefficient is efficient for tree-ring analyses, once the samples vary in length and periods. Based on the dissimilarity matrix a constrained hierarchical clustering (where the agglomeration method is the Constrained Incremental Sums of Squares - CONISS) was performed with restricted clusters in order of the years (Gordon & Birks, 1972), the function was implemented on the rioja R package (Juggins, 2017).

To access the changes over time, we broke the data into three periods of the same length (33 years) for each area, two before (P1: 1917-1949 and P2: 1950-1982) and one after (P3: 1983-2015) the Balbina Dam construction which started in 1983 for trees from the USDR. We then verified if there were differences in growth between periods within the area and if the same patterns persist for trees from the JNP. Finally, we compared the periods between areas, considering in the JNP the intensification on the hydrological cycle, which started around the same period (Barichivich et al., 2018; Gloor et al., 2013) as the Uatumã river damming. For comparing the MDI between equal periods inside each area and between areas, we first performed Shapiro-Wilk normality tests to proceed with the Wilcoxon rank-sum test for non-parametric datasets or t-tests for parametric ones (R Core Team, 2018).

#### Determination of the date of death using ^14^C dating

We dated the year of death for 41 trees (only those with intact outermost rings), 13 from JNP e 28 from USDR. The year of death was determined by comparing the radiocarbon (^14^C) content of cellulose extracted from the outermost (final) ring of each sample with the atmospheric record of ^14^C in the southern hemisphere. Cellulose was isolated from bulk wood using the Jayme-Vise method (Green, 1963; Axel Steinhof, 2013) in the ^14^C Analysis Facility at the Max-Planck Institute for Biogeochemistry in Jena, Germany (A. Steinhof, Altenburg, & Machts, 2017). Radiocarbon measurements were made on graphite targets using an accelerator mass spectrometer (AMS), reporting the results as Fraction Modern, or the ratio of ^14^C/^12^C in the sample (corrected to a ^13^C of −25, compared to that of a standard representing the preindustrial atmosphere). Calibrated ages were determined from the Fraction Modern of cellulose using the OxCal online software v4.3 (Ramsey, 2001; Showman, Wordsworth, Merlis, & Kaspi, 2013) and considering the concentration of radioactive carbon in the modern standard (post-1950 period /Bomb 13 SH Zone 3 curve) (Hua, Barbetti, & Rakowski, 2013).

For deaths occurring in the pre-bomb interval between ~1650 and 1950 AD, when errors associated with radiocarbon dating are large due to measurement precision combined with atmospheric ^14^C changes (Suess-effect) (Hua et al., 2013), or cases of more than one possible year or a range of years indicated by Oxcal after the bomb period, we used ring counting to estimate the most plausible death date. These dates were compared with the history of disturbances induced by the hydroelectric dam (for the USDR trees) and severe hydroclimatic events (both areas) associated with large-scale climate anomalies such as the *El Niño*-Southern Oscillation (ENSO) using the Southern Oscillation Index (SOI) data (CRU, 2019; Ropelewski & Jones, 1987) to understand the cause of their death.

## Results

The mean DBH and age were similar for both areas, being 43.9 cm (min-max: 16.5-116 cm) and 210 yrs (60-466 yrs) for the JNP trees and 45.9 cm (12.1-175.07 cm) and 213 yrs (30-498 yrs) for USDR trees. Consequently, the average MDI values showed little but significant difference (*p* = 0.01, *t*-test), between both sites (2.04 cm in JNP and 2.28 cm in USDR) (Table 1). The ages and sizes of hollows were higher for tree from the USDR (104 yrs with 22.4 cm) compared to the JNP (77 yrs with 17.2 cm).

**Table 1.** Quantitative results of the studied populations of Eschweilera tenuifolia at the Jaú National Park (JNP) and Uatumã Sustainable Development Reserve (USDR) comparing mean, standard deviation (SD), minimum and maximum values of diameter at breast height (DBH), the diameter and age of the hollow (HolAge), estimated age and mean diameter increment (MDI). Asterisks indicate significant differences using a t-test (p <0.05).

|  | Area | DBH (cm) | Hollow (cm) | HolAge (yr) | Total Age (yrs) | MDI (mm) |
| --- | --- | --- | --- | --- | --- | --- |
| **Num.** | JNP | 52 | 38 | 38 | 52 | 52 |
| **Ind.** | USDR | 91 | 57 | 57 | 90 | 91 |
| **Mean** | JNP | 43.9 ± 21.7 | 17.2 ± 13.8 | 77 ± 61 | 210 ± 103 | 2.04 ± 0.39* |
| ± SD | USDR | 45.9 ± 24.6 | 22.4 ± 23.0 | 104 ± 76 | 213 ± 103 | 2.28 ± 0.69* |
| Min. | JNP | 16.5 | 2.3 | 17 | 60 | 1.29 |
|  | USDR | 12.1 | 4.0 | 24 | 30 | 1.10 |
| Max. | JNP | 116.0 | 63.2 | 287 | 466 | 2.97 |
|  | USDR | 93.6 (175.2) | 60.0 | 272 | 498 | 4.88 |

\*: the individual with the largest DBH was not considered for estimated tree age, as the age estimate of the large hollow had methodological restrictions.

The age-DBH model for living and dead, hollow and non-hollow trees demonstrates that the growth trajectory is similar in both areas, although the trees at the USDR presented a higher variability in diameter growth (Figure S6). At both sites, the cumulative DBH is highly correlated to the total ages (*r*=0.97 and R^2^= 0.95 for JNP and *r*=0.96 and R^2^= 0.92 for USDR).

The MDI-pattern to pass through 5-cm diameter classes is well defined for the population from the JNP, but not in the USRD that presented a greater variation in values. The MDI was significantly different (*p* = 0.01, *t-test*) between areas but not between living and dead trees for each area. In both systems, the species showed the lowest increment rates in diameter in the first class (0 to 5 cm) with no significant difference between living (JNP: 1.67 mm year^−1^; USDR: 1.50 mm year^−1^) and dead (JNP: 1.94 mm year^−1^; USDR: 1.38 mm year^−1^), meaning that in this class trees spent more time to accumulate 5 cm than in any the subsequent classes. Maximum MDIs were attained in the fifth diameter class in the USDR (2.81 mm year^−1^ for living trees) and already in the third class on the JNP (2.99 mm year^−1^ for living trees). After 55 cm in diameter, the increment rates tended to increase for dead trees from the JNP, but it is not well defined for the population from the USDR (Figure 3). The notched boxplots are tests of differences per class between living and dead individuals within each area. At the JNP there was a difference in the class 15-20 cm, while in the USDR differences were observed. Comparing the dead trees between both areas indicated significant differences from the class 15-20 to 25-30 cm, however, the same was not observed for living trees.

Analyzing the growth ratio preceding death (*g*_*r*_), we identified for 13 individuals (62%) from the JNP at least one year with *g*_*r*_<1.0 before death. At the USDR 15 trees (52%) showed this pattern. This means that the other 38% of the JNP and 48% of the USDR died abruptly, without an apparent decline in preceding years. The number of consecutive years with *g*_*r*_<1.0 before death, however, varied between both study sites. At the JNP they spent the five-fold time to die in comparison to the USDR (Figure 5).

**Figure 5.**
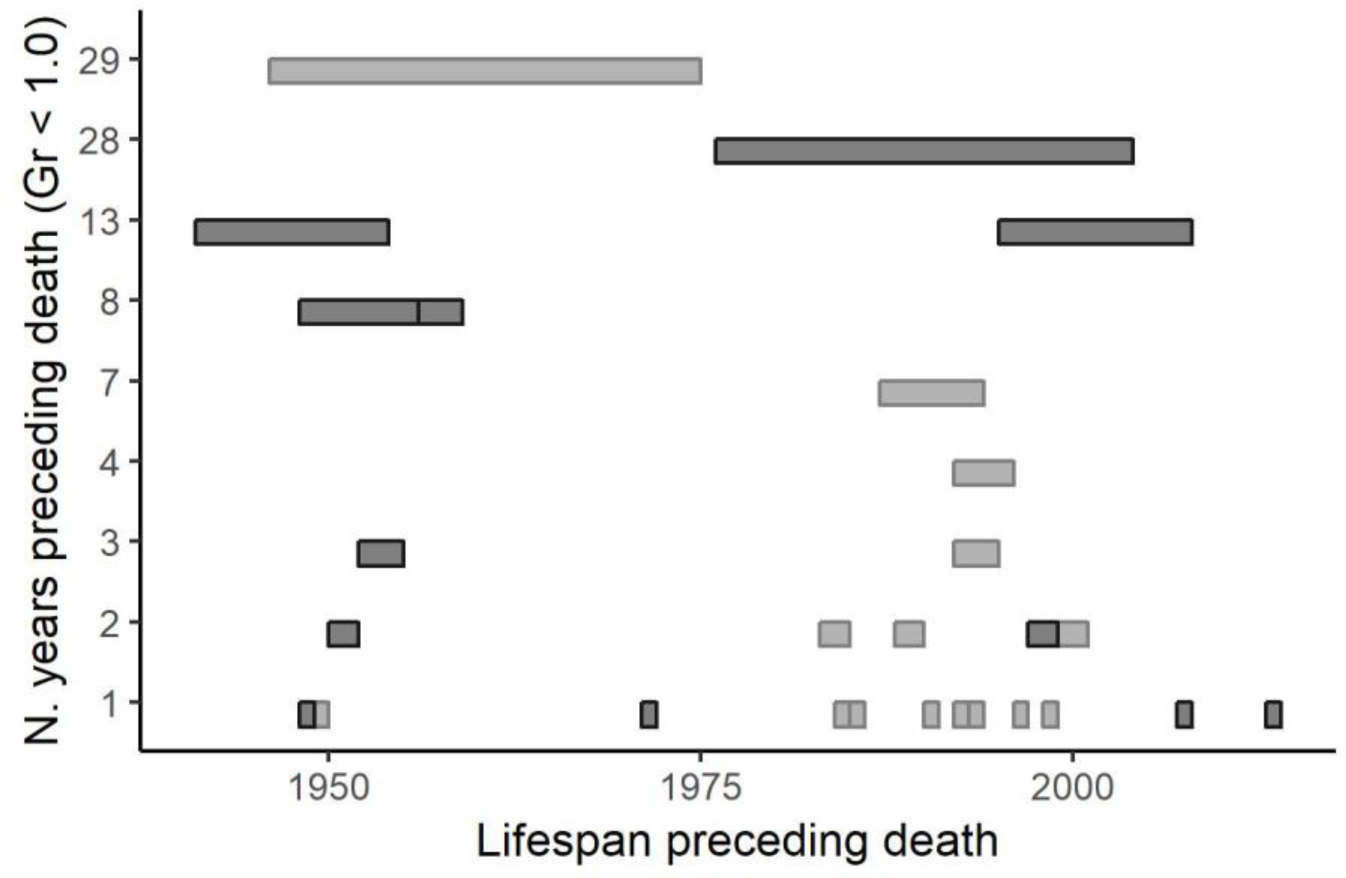
The period preceding the death of each individual of Eschweilera tenuifolia with gr <1.0 for the Jaú National Park (JNP; dark grey) and the Uatumã Sustainable Development Reserve (USDR; light grey).

Using cluster analysis, living trees from the USDR were grouped in two distinct clusters (Figure 6a) split by the years 1981-1982, while in the JNP living trees were separated by the period 1975-76, also split into two main blocks (Figure 6a). In each block, subdivisions can be noticed. At the USDR site, 17 individuals (18.7% of all trees) presented release and suppression events which occurred more frequently at closed mixed forests (58.3%), compared to the opened monodominant formation in lakes (12.7%) (Table S1). Among trees from lakes dead trees (17.2%) presented more frequently events of growth changes than living trees (10.0%). Eight of the 10 living trees with growth suppression presented the synchronized in the post-dam-period indicating exogenous disturbances (Figure 6a) although synchronous release events happened in the USDR also before the dam construction. This pattern was quite different for trees from the JNP where only 11.5% of all analyzed trees showed significant growth changes. Those of living trees were only observed in closed mixed forests where 30.8% of the analyzed trees presented not synchronized release and/or suppression events indicating endogenous disturbances (inter-tree competition). Only 9.5% of the dead trees from macrohabitat at lakes showed growth changes.

**Figure 6.**
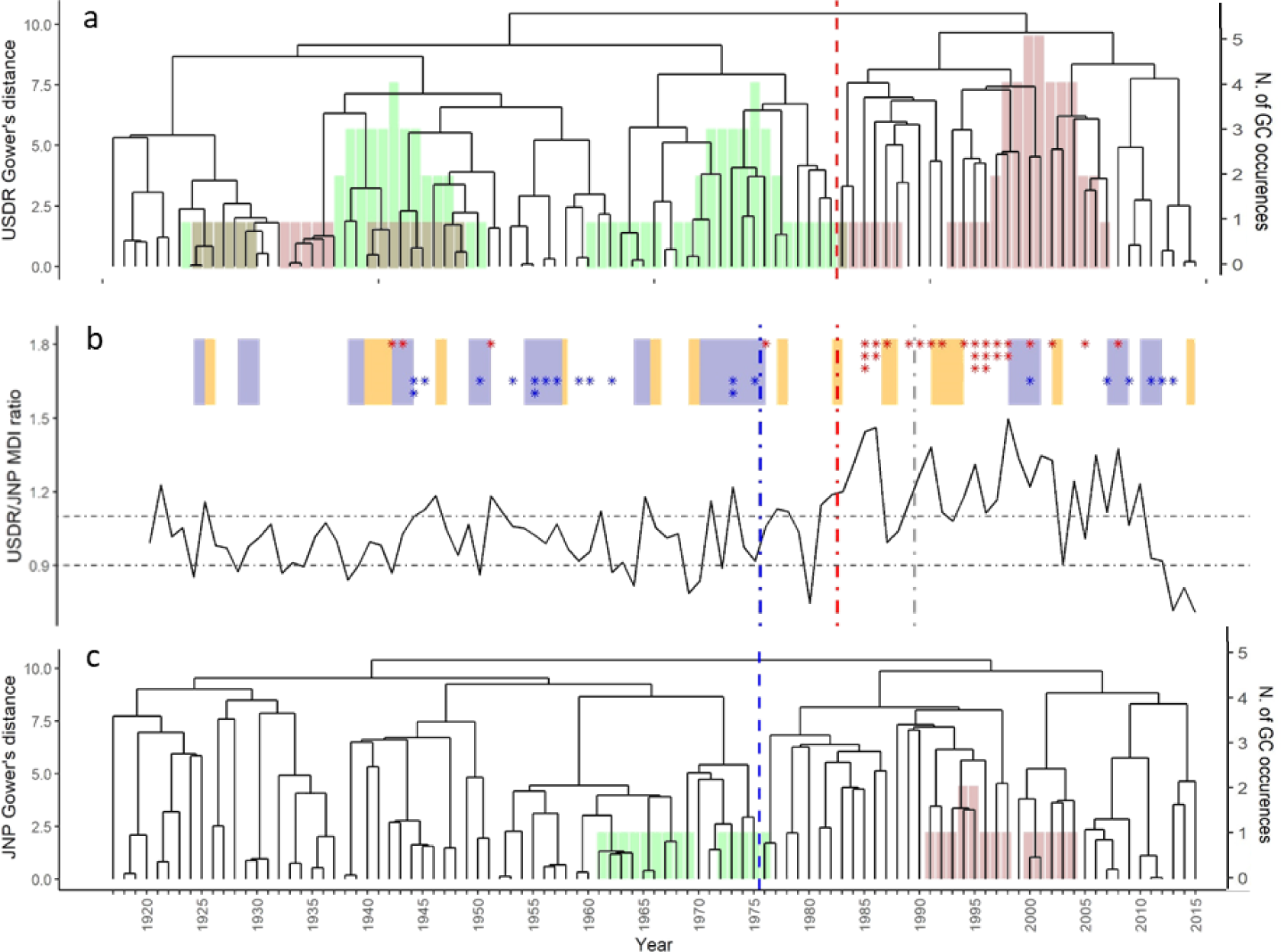
The panels 6a and 6c represents the cluster analysis for (a) Uatumã Sustainable Development Reserve (USDR) and (c) Jaú National Park (JNP). The vertical lines represent the split-year of the growth in the cluster analysis for USDR (red) and JNP (blue). The diameter growth changes (%) of trees presenting abnormal growth patterns at (a) USDR and (c) JNP are shown in the bars, being suppression (<-50%, in pink) and release events (>100%, in green). Figure 6b shows the mean diameter increment (MDI) ratio between USDR and JNP (dot-slash horizontal lines are showing 10% thresholds). The asterisks highlight the estimated year of death for each individual from USDR (in red) and JNP (in blue) and the colored rectangles indicate El Niño (yellow) and La Niña (blue) events.

For dead trees, the modern radiocarbon analysis showed that 87% of the mortality at the USDR occurred after the begin of the implementation of the Balbina dam, while mortality occurred during different periods at the JNP mostly associated with *La Niña* years (Figure 6b). After assuming that ontogenetic effects on MDI were overcome by sampling trees with different sizes, we compared for each study site the MDI since 1919 for equal intervals based on the 33-years post-dam period, where P3: 1983-2015; P2: 1950-1982; P1: 1917-1949. The population growing at the USDR showed no differences for the pre-dam periods P1 and P2, but significant differences between pre-dam periods and the post-dam period P3. For the JNP-population, no differences between any periods were observed. Comparing the populations between USDR and JNP, we found significant differences for the period P3 which reflect the period of intensification of the hydrological cycle induced by the implementation of the Balbina dam leading to increased minimum water levels (USDR) (Assahira *et al*., 2017) and by increasing maximum water levels at the JNP associated by oceanic forcing (Barichivich *et al*., 2018) (Table S2).

To have a clear idea of the magnitude of differences in the MDI of living trees between both areas, we calculated the MDI ratio (USDR/JNP; values >1 indicate higher diameter growth at the USDR) (Figure 6b). Tree growth was considerably lower at the USDR in the period from 1960-1965. After 1985 the population of the USDR showed much faster diameter growth compared to the JNP with an abrupt steep decline starting around 2002 resulting in a growth ration below 0.9 after 2012 (Figure 6b).

## Discussion

The tree species *E. tenuifolia* growing at harsh environmental conditions in the highly flooded and oligotrophic igapó reach high tree ages. The oldest living tree was found at the USDR with a DBH of 175 cm, but its age had to be estimated by a linear regression due to the large hollow trunk section. Considering the individual MDI of 2.15 mm this tree was estimated to be over 800 years old, although the slow growth pattern in the first classes (1.50 mm for USDR living trees) was not accounted, what could substantially increase its age. This estimate is much higher than former indicated maximum ages for tree species in this ecosystem with 502 years for a *M. acaciifolium* individual from the *igapó* forests of the Amanã Lake, Central Amazonia, using linear models to estimate the hollow part of the trunk (Schöngart et al., 2005).

Retrospective analyses performed in this study combining tree-ring analyses, radiocarbon dating, and analysis of long-term hydrographic data allows us to get more insights into the ecological processes in this oligotrophic *igapó* characterized by low dynamical processes and the growth dynamics of the populations. Growth anomalies seem to be more frequent at the disturbed *igapó* site compared to the pristine conditions (JNP) but are more restricted to the post-dam conditions at the USDR. Although MDI was slightly higher at the USDR, similar trends in growth rates were recorded for both study areas, which demonstrates an ontogenetical resemblance. However, analyzing the growth patterns shows some interesting differences among the study sites. Tree growth at the JNP was less variable compared to the USDR site and only some individuals indicated major changes in diameter growth leading to isolated release and suppression events. Interestingly, these events were only recorded for trees growing in closed-canopy formations under high inter-tree competition with some other highly flood-adapted tree species, such as *Burdachia prismatocarpa* A. Juss., *Buchenavia ochroprumna* Eichler, *Amanoa oblongifolia* Müll. Arg. and *M. acaciifolium* (Aguiar, 2015; Leandro V Ferreira, 1997) possible resulting from endogenous disturbances (competition). Release events occurred before dam construction at USDR, isolated or synchronous among individuals indicating endogenous, but possibly also exogenous disturbances such as the periods 1935-1947 and1968-1976. Different responses to rainfall anomalies in the watersheds and consequently hydrological regimes which are thought to be the main trigger of tree-growth variability in the Central Amazonian floodplains (Schöngart et al., 2004, 2005). The available river discharge records from the National Agency of Waters (ANA) from 1931 to 2015 (ONS, 2016) show a low discharge in 1937 (Figure 3S) which could have caused the first release, but we cannot explain the second one by hidro-climatic events. Synchronous suppression events at the USDR site are mainly observed during the period of operation of the Balbina dam, especially after 2000 (Figure 6c) which corresponds to the begin of a significant increase of the low-water regime of the Uatumã River during consecutive years (Assahira et al., 2017).

At the JNP tree mortality occurred during different periods mainly during the periods 1946-1975 and 1999-2014. Interestingly, both periods coincide with cold phases of the Interdecadal Pacific Oscillation (IPO), a decadal climate variability across the Pacific Ocean modulating interannual variability of the ENSO (Salinger *et al*., 2016). Barichivich *et al.* (2018) suggest that the IPO is modulating the hydrological regime in the Central Amazon region, increasing maximum water levels during cold phases. The majority of tree’s death (>70%) coincide with *La Niña* years (cold ENSO phases) which causes enhanced annual maximum and minimum water levels and prolonged flooding periods (Schöngart *et al*., 2004; Schöngart and Junk, 2007, in press) which possibly is the main trigger of mortality pattern observed at the JNP. Extreme hydroclimatic events occurred between 1971 and 1975 resulting in the extinction of the terrestrial phase at the low floodplain topographies in Central Amazonia for several years (Piedade *et al*., 2013). Enhanced mortality of shrub and tree species growing at these macrohabitats was observed which is consistent with our findings from the USDR which can be considered as a field experiment (Assahira *et al*., 2017; Resende *et al*., 2019). Although these hydroclimatic disturbances were not severe enough to cause massive mortality of the populations growing under pristine conditions at the JNP, we suggest that these events are the main trigger of tree mortality, sporadically killing small cohorts especially in cold phases of ENSO and IPO which strongly modulate the hydrological regime in the Central Amazonian floodplains (Barichivich *et al*., 2018; Schöngart and Junk, 2007, in press) (Figure 6b). We also observe a major change in the MDI patterns after the severe hydroclimatic event from the 1970s at the JNP extending into the period of the intensification of the hydrological cycle (Gloor *et al*., 2013, 2015; Barichivich *et al*., 2018).

At the USDR considerable changes in the MDI-pattern occurred around the begin of the Balbina dam construction period (1983-1987). In October 1987 the dam was closed for filling the large reservoir to initiate the hydroelectric power generation in February 1989 (Fearnside, 1990a). This reduced the downstream discharge and consequently increased the terrestrial phase in the *igapó* floodplains (Neves et al., 2019).. The prolonged terrestrial phase might have favored diameter growth increases during this period (Figure 6b) as suggested by Schöngart *et al*. (2004, 2005). Possibly these trees growing at low elevations possibly had groundwater access (M. T. F. Piedade et al., 2013). However, with begin of the Balbina dam construction we also observed massive tree mortality of *E. tenuifolia*. Of all analyzed dead trees, 87% died after the begin of the construction of the Balbina dam; one third of those died during the period of implementation (1983-1988) and the other two thirds after the beginning of the Balbina dam operation in 1989. We have two hypotheses for the observed mortality during the period of dam implementation both associated to extreme hydroclimatic events induced by *El Niño* conditions which occurred during this period in the years 1982/1983 and 1986-1988.

Mainly during strong *El Niño* events, severe hydroclimatic droughts in the igapó floodplains may turn these forests extremely vulnerable to wildfires (Flores et al., 2017; Flores, Piedade, & Nelson, 2014; Schöngart, Wittmann, Junk, & Piedade, 2017). The air humidity in *igapó* forests on lower topographies is considerably lower compared to the stratified canopy of adjacent upland forests (Resende *et al*., 2014; Almeida *et al*., 2016). A large amount of fine fuels accumulates on the forest floor due to the slow decomposition of fine litter during the anaerobic conditions caused by long inundations and a dense root mat formed close to the soil surface (dos Santos and Nelson, 2013). The temperature during these extreme events is increased, and the relative air humidity lowered due to the reduced rainfall associated with *El Niño* (Aragão *et al*., 2018; Marengo *et al*., 2018) also lowering river discharge. In synergy with the Balbina dam implementation this might have create conditions for wildfires which also might affect the macrohabitats dominated by *Eschweilera tenuifolia* causing mortality. This hypothesis is sustained by identifying young cohorts of fast-growing pioneer species in the *igapó* forests downstream of the Balbina dam which possibly established after wildfires occurring in this period (Neves *et al*., 2019).

Another explanation for the mortality mechanisms can be linked to hydraulic failure or depletion in carbon reserves (Cailleret et al., 2017; Sevanto, Mcdowell, Dickman, Pangle, & Pockman, 2014), but the trigger behind the depletion of carbon in healthy trees are still poorly understood (Bucci, Goldstein, Scholz, & Meinzer, 2016). Hydraulic failure is mainly caused by xylem embolism; however, it strongly depends on the hydraulic architecture associated with wood anatomical features (vessel size and distribution, pit membrane thickness between adjacent vessels) (Cailleret et al., 2017; Zeppel, Adams, & Anderegg, 2011). Losses of water potential (ψ), derived from xylem vulnerability curves under controlled conditions, are suggested to lead to mortality at ψ50 or ψ88 (loss of 50% or 88%) (Li et al., 2016; Urli et al., 2013). When the net carbon balance becomes negative, non-structural carbohydrates (NSC) reserves sustain vital processes such as respiration (Hartmann, Adams, et al., 2018). The water deficit during *El Niño* events in the Central Amazon in combination with higher temperatures and increasing vapor pressure deficits (VPD) might cause a negative net carbon balance. During the period of flooding, many tree species switch the metabolism from aerobic to anaerobic pathways to maintain basic functional processes, mobilizing NSC stored during the terrestrial phase (Parolin et al., 2004; Schlüter & Furch, 1992). When the NSC storages are depleted after inundations, floodplain trees must renew those by photosynthesis and mobilization, but the occurrence of droughts as a consequence of reduced rainfall, high temperatures and increased VPD might limit the NSC recover during the terrestrial phase which is the growing season in the Amazonian floodplains (Schöngart *et al*., 2002). A recent study performing stable carbon (δ^13^C) and oxygen (δ^18^O) isotope analyses of tree rings at an intra-annual resolution suggest for *M. acaciifolium* that local climate conditions (temperature, cloud cover, precipitation, dry season length) may reach increasingly stressful levels affecting growth performance by increasing leaf-to-air VPD leading to stomatal closure and leaf water enrichment (Cintra et al., 2019). In consequence, the tree might die due to carbon starvation.

Interestingly, the mortality of the trees at the USDR during the period of dam implementation occurred abruptly for the majority of analyzed trees, without a long growth decline (*g*_*r*_) in preceding years. Under pristine conditions at the JNP no *E. tenuifolia* tree died during this period. Furthermore, dead trees from the JNP presented a much longer growth rate declines before death, suggesting different triggers and causes of mortality between both sites (Figure 5). One hypothesis (wildfires) does exclude the other (hydraulic failure/carbon starvation) as the cause for sudden tree mortality at the USDR. However, future studies are necessary to get more insights for the causes of mortality, such as mapping areas and periods of wildfires affecting *igapó* forests in the USDR and achieve measurements of the ecophysiological performances combined with analyses of functional leave and wood traits as well as anatomical features related to the hydraulic conduction (Fichtler & Worbes, 2012; Hartmann, Moura, et al., 2018; G. B. Mori, Schietti, Poorter, & Piedade, 2019; Zeppel et al., 2011). Moreover, it is necessary to screen the mechanism of adaptations that this species developed to tolerate the long periodic inundation.

During the period of dam operation starting in February 1989, the scenario changed dramatically leading to an increase of the low-water regime in the period 1994-1997 causing the extinction of the terrestrial phase inducing the second wave of mortality of *E. tenuifolia* trees, possibly to prolonged anoxic conditions. After the strong *El Niño* event in 1997/1998, causing again increased diameter growth (and also tree mortality), permanent flooding condition initiated around 2000 causing permanent inundations for over 10 years which resulted into mass mortality of *M. acaciifolium* growing on slightly higher elevations in the *igapó* (Assahira *et al*., 2017). During this period, we observe a strong decline in MDI of the living trees of *E. tenuifolia* especially in the period after 2002 and also mortality of some trees. Nowadays, dead stands comprise 12% of the area of floodplain forests (about 1,100 ha) up to 120 km downstream of the dam, mainly affecting macrohabitats dominated by *E. tenuifolia* (Resende *et al*., 2019). The declining growth rates of still-living trees suggest that a significant portion of this population will not overcome the persistent disturbance suffering mortality in the next future, as postulated by Resende *et al.* (2019).

Taking into account the complex disturbance regimes that influence mortality patterns in the Amazonian floodplains, which cover over 750.000 km^2^ in the Amazon basin (Junk et al 2011) only a few studies have shown the effects of river damming and fires on *igapó* forests (Almeida et al., 2016; Assahira et al., 2017; Flores et al., 2017, 2014; Nelson, 2001; Angélica F. Resende et al., 2014; Angélica Faria de Resende et al., 2019; Schöngart, Wittmann, et al., 2017). In upland forests (*terra firme*), for instance, many studies highlight the effects of different disturbances agents on forest structure, biodiversity, carbon balance, and tree mortality. Natural tree mortality (de Toledo, Magnusson, Castilho, & Nascimento, 2012), extreme climatic events such as windstorms (Negrón-Juárez et al., 2010), droughts (Aleixo et al., 2019; Phillips et al., 2009), wildfires (Barlow & Peres, 2008; Brando et al., 2014) and other causes related to anthropogenic factors such as forest fragmentation (Williamson et al., 2000) are known disturbance agents leading to enhanced tree mortality and losses of ecosystem services. Additionally, the complex environmental interactions in an area that culminate in the mortality of trees over time is one of the less-studied processes in ecology (Franklin, Shugart, & Harmon, 1987; Villalba & Veblen, 1998), even though it is of fundamental importance for the understanding of forest ecology and dynamics (Johnson & Fryer, 1989; Oliver & Larson, 1996; Veblen, 1986). In comparison to the *terra firme* environments, the mortality triggers in the *igapó* floodplains may be different as it also was observed for tree growth (Schöngart *et al*., 2004, 2010; Cintra *et al*., 2019).

We conclude that alterations in the hydrological regimes caused by climate and land-use changes and synergies between those are affecting the macrohabitats dominated by *E. tenuifolia*. This possibly endemic tree species from the *igapó* is extremely sensitive and vulnerable to both, long- and short-lasting abnormal flooding associated to extreme hydroclimatic or anthropogenic disturbances. Massive mortality of these populations are leading to irreplaceable losses of very special macrohabitats and ecosystem services (Wolfgang J. Junk et al., 2018). Hydroelectric dams cause abrupt and massive changes in the floodplains downstream of the reservoir (Timpe and Kaplan 2017) which affect tree growth and mortality especially at these macrohabitats (Assahira *et al*., 2017; Resende *et al*. 2019). This results in losses of macrohabitat, genetic resources of one of the best flood-adapted tree species providing important ecosystem services. Under natural conditions tree mortality at these macrohabitats of the *igapó* seems to be triggered during cold phases of large-scale oscillations originating from the Pacific Ocean at high (ENSO) and low (IPO) frequencies, but possibly enhance tree growth and cohort establishment during warm phases of ENSO (*El Niño*) and IPO (Schöngart *et al*., 2004, 2005). However, the increase of the mean water level of 1 m during a period of 116 years (Barichivich *et al*., 2018), probably will lead to a slowly ongoing extinction of these macrohabitats and force trees to occupy macrohabitats on higher floodplain elevations.

## Acknowledgments

We thank the National Institute of Amazonian Research (INPA), the Laboratory of Dendroecology (Casa 20), the PELD/MAUA group, INCT ADAPTA, the Botany Post-graduation, the Jaú National Park (from ICMBio), the Uatumã Sustainable Development Reserve (SEMA/DEMUC) and ATTO (Amazon Tall Tower Observatory) project for the infrastructure and technical support. We acknowledge support from the Brazilian Council for Scientific and Technologic Development (CNPq) for financing the project and providing the Ph.D. fellowship for A.F. Resende, the LBA-CNPq grant (MCTI/CNPq/FNDCT nº 68/2013 - Large-Scale Biosphere-Atmosphere Program in the Amazon - LBA), PELD-MAUA project (403792/2012-6 - MCTI/CNPq/FAPs Nº 34/2012 and 441590/2016-0 - CNPq/CAPES/FAPS/BC-Fundo Newton). We must thank also for the support from IBAMA, SISbio, SISgen, ICMBio, and DEMUC/SEMA, for the licenses for collecting and transporting the wood samples (licenses 51133-1/2015 and 61/2015 - 28/2018 from SISBio and DEMUC/SEMA, respectively). We also thank Reginaldo, Victor Lery, Eduardo Rios, Jaime, Alberto Peixoto, Josué, Domingos, Celso Rabelo, Jekiston Andrade, Ketlen Fernanda, Anderson, and Gildo Feitoza for their support during fieldwork. This manuscript preprint is part of A. F. Resende Ph.D.-thesis accessible at INPA digital library. We thank Tim Vincent and Adrian Barnett for the English revisions.

## References

Aguiar, D. P. P. (2015). Influência dos fatores hidro-edáficos na diversidade, composição florística e estrutua da comunidade arbórea de igapó no Parque Nacional do Jaú, Amazônia Central.

Aleixo, I., Norris, D., Hemerik, L., Barbosa, A., Prata, E., Costa, F., & Poorter, L. (2019). Amazonian rainforest tree mortality driven by climate and functional traits. Nature Climate Change. https://doi.org/10.1038/s41558-019-0458-0

Almeida, D. R. A. de, Nelson, B. W., Schietti, J., Gorgens, E. B., Resende, A. F., Stark, S. C., & Valbuena, R. (2016). Contrasting fire damage and fire susceptibility between seasonally flooded forest and upland forest in the Central Amazon using portable profiling LiDAR. Remote Sensing of Environment, 184, 153–160. https://doi.org/10.1016/j.rse.2016.06.017

Andreae, M. O., Acevedo, O. C., Araùjo, A., Artaxo, P., Barbosa, C. G. G., Barbosa, H. M. J., … Yáñez-Serrano, A. M. (2015). The Amazon Tall Tower Observatory (ATTO): Overview of pilot measurements on ecosystem ecology, meteorology, trace gases, and aerosols. Atmospheric Chemistry and Physics, 15(18), 10723–10776. https://doi.org/10.5194/acp-15-10723-2015

Armstrong, W., & Drew, M. C. (2002). Root growth and metabolism under oxygen deficiency. Plant Roots: The Hidden Half, 729–761. https://doi.org/10.1016/S0079-8169(04)80088-6

Artaxo, P. (2019). Working together for Amazonia. Science, 363(6425), 323–323. https://doi.org/10.1126/science.aaw6986

Assahira, C., Piedade, M. T. F., Trumbore, S. E., Wittmann, F., Cintra, B. B. L., Batista, E. S., … Batista, E. S. (2017). Tree mortality of a flood-adapted species in response of hydrographic changes caused by an Amazonian river dam. Forest Ecology and Management, 396, 113–123. https://doi.org/10.1016/j.foreco.2017.04.016

Baker, P. J., Bunyavejchewin, S., Oliver, C. D., & Ashton, P. S. (2005). Disturbance history and historical stand dynamics of a seasonal tropical forest in western Thailand. Ecological Monographs, 75(3), 317–343. https://doi.org/10.1890/04-0488

Barichivich, J., Gloor, E., Peylin, P., Brienen, R. J. W., Schöngart, J., Espinoza, J. C., … Schöngart, J. (2018). Recent intensification of Amazon flooding extremes driven by strengthened Walker circulation. Science Advances, 4(9), eaat8785. https://doi.org/10.1126/sciadv.aat8785

Barlow, J., & Peres, C. A. (2008). Fire-mediated dieback and compositional cascade in an Amazonian forest. Philosophical Transactions of the Royal Society B: Biological Sciences, 363(1498), 1787–1794. https://doi.org/10.1098/rstb.2007.0013

Barnett, A. A., Borges, S. H., Shapley, R. L., Castilho, C. V. de, Neri, F. M., & Shapley, R. L. (2002). Primates of the Jaú National Park, Amazonas, Brazil. Neotropical Primates, 10(2), 65–70.

Barnett, A. A., Bowler, M., Bezerra, B. M., & Defler, T. R. (2013). Ecology and behavior of uacaris (genus Cacajao). In Evolutionary Biology and Conservation of Titis, Sakis and Uacaris (pp. 151–172). https://doi.org/10.1017/cbo9781139034210.020

Barnett, A. A., Castilho, C. V. De, Shapley, R. L., & Anicácio, A. (2005). Diet, Habitat Selection and Natural History of Cacajao melanocephalus ouakary in Jaú National Park, Brazil1. International Journal of Primatology, 26(4), 949–969. https://doi.org/10.1007/s10764-005-5331-5

Batista, E. S., & Schöngart, J. (2018). Dendroecology of Macrolobium acaciifolium (Fabaceae) in Central Amazonian floodplain forests. Acta Amazonica, 48(4), 311–320. https://doi.org/10.1590/1809-4392201800302

Bauer, D. F. (1972). Constructing confidence sets using rank statistics. Journal of the American Statistical Association, 67(339), 687–690. https://doi.org/10.1080/01621459.1972.10481279

Beissner, A., Genske, R., Prall, M., & Weinholz, P. (2002). Xact Version 7.22b - SciLab GmbH. Germany.

Bowman, D. M. J. S., Brienen, R. J. W., Gloor, E., Phillips, O. L., & Prior, L. D. Detecting trends in tree growth: Not so simple., 18 Trends in Plant Science § (2013).

Brando, P. M., Balch, J. K., Nepstad, D. C., Morton, D. C., Putz, F. E., Coe, M. T., … Soares-Filho, B. S. (2014). Abrupt increases in Amazonian tree mortality due to drought-fire interactions. Proceedings of the National Academy of Sciences, 111(17), 6347–6352. https://doi.org/10.1073/pnas.1305499111

Brienen, R. J. W., & Zuidema, P. A. (2006). Lifetime growth patterns and ages of Bolivian rain forest trees obtained by tree ring analysis. Journal of Ecology, 94(2), 481–493. https://doi.org/10.1111/j.1365-2745.2005.01080.x

Brienen, R. J. W., Zuidema, P. A., & Martínez-Ramos, M. (2010). Attaining the canopy in dry and moist tropical forests: Strong differences in tree growth trajectories reflect variation in growing conditions. Oecologia, 163(2), 485–496. https://doi.org/10.1007/s00442-009-1540-5

Bucci, S. J., Goldstein, G., Scholz, F. G., & Meinzer, F. C. (2016). Physiological Significance of Hydraulic Segmentation, Nocturnal Transpiration and Capacitance in Tropical Trees: Paradigms Revisited. https://doi.org/10.1007/978-3-319-27422-5_9

Cailleret, M., Jansen, S., Robert, E. M. R., Desoto, L., Aakala, T., Antos, J. A., … Martínez-Vilalta, J. (2017). A synthesis of radial growth patterns preceding tree mortality. Global Change Biology, 23(4), 1675–1690. https://doi.org/10.1111/gcb.13535

Castello, L., Mcgrath, D. G., Hess, L. L., Coe, M. T., Lefebvre, P. A., Petry, P., … Arantes, C. C. (2013). The vulnerability of Amazon freshwater ecosystems. Conservation Letters, 6(4), 217–229. https://doi.org/10.1111/conl.12008

Cintra, B. B. L., Gloor, M., Boom, A., Schöngart, J., Locosselli, G. M., & Brienen, R. (2019). Contrasting controls on tree ring isotope variation for Amazon floodplain and terra firme trees. Tree Physiology, 1–16. https://doi.org/10.1093/treephys/tpz009

Cook, E., & Kairiukstis, L. (1990). Methods of dendrochronology (E. R. Cook & L. A. Kairiukstis, Eds.). https://doi.org/10.1016/0048-9697(91)90076-q

Crawford, R. M. M., & Braendle, R. (2007). Oxygen deprivation stress in a changing environment. Journal of Experimental Botany, 47(2), 145–159. https://doi.org/10.1093/jxb/47.2.145

CRU, C. R. U. (2019). Southern Oscillation Index (SOI) Archives. Retrieved December 9, 2019, from https://crudata.uea.ac.uk/cru/data/soi/

Da Fonseca Júnior, S. F., Piedade, M. T. F., & Schöngart, J. (2009). Wood growth of Tabebuia barbata (E. Mey.) Sandwith (Bignoniaceae) and Vatairea guianensis Aubl. (Fabaceae) in Central Amazonian black-water (igapó) and white-water (várzea) floodplain forests. Trees - Structure and Function, 23(1), 127–134. https://doi.org/10.1007/s00468-008-0261-4

Davidson, E. A., De Araüjo, A. C., Artaxo, P., Balch, J. K., Brown, I. F., Mercedes, M. M., … Wofsy, S. C. (2012). The Amazon basin in transition. Nature, 481(7381), 321–328. https://doi.org/10.1038/nature10717

de Toledo, J. J., Magnusson, W. E., Castilho, C. V, & Nascimento, H. E. M. (2012). Tree mode of death in Central Amazonia: Effects of soil and topography on tree mortality associated with storm disturbances. Forest Ecology and Management, 263, 253–261.

Fearnside, P. M. (1990a). Balbina: lições trágicas na Amazônia. Ciência Hoje, 11(64), 34–40.

Fearnside, P. M. (1990b). Environmental Destruction in the Brazilian Amazon. The Future of Amazonia, (January 1990), 179–225. https://doi.org/10.1007/978-1-349-21068-8_8

Fearnside, P. M. (2005). Deforestation in Brazilian Amazonia: History, rates, and consequences. Conservation Biology, 19(3), 680–688. https://doi.org/10.1111/j.1523-1739.2005.00697.x

Ferreira, L. V. (2000). Effects of flooding duration on species richness, floristic composition and forest structure in river margin habitat in Amazonian blackwater floodplain forests: implications for future design of protected areas. Biodiversity and Conservation, 9(1), 1–14. https://doi.org/10.1023/A:1008989811637

Ferreira, Lv, & Prance, G. T. (1998). Structure and species richness of low-diversity floodplain forest on the Rio Tapajós, Eastern Amazonia, Brazil. Biodiversity and Conservation, 7, 585–596. https://doi.org/10.1023/A:1008848200441

Ferreira, Leandro V. (1997). Effects of the duration of flooding on species richness and floristic composition in three hectares in the Jau National Park in floodplain forests in central Amazonia. Biodiversity & Conservation, 6(10), 1353–1363. https://doi.org/10.1023/A:1018385529531

Fichtler, E., & Worbes, M. (2012). Wood anatomical variables in tropical trees and their relation to site conditions and individual tree morphology. IAWA Journal, 33(2), 119–140. https://doi.org/10.1163/22941932-90000084

Flores, B. M., Holmgren, M., Xu, C., van Nes, E. H., Jakovac, C. C., Mesquita, R. C. G., & Scheffer, M. (2017). Floodplains as an Achilles’ heel of Amazonian forest resilience. Proceedings of the National Academy of Sciences, 114(17), 4442–4446. https://doi.org/10.1073/pnas.1617988114

Flores, B. M., Piedade, M. T. F., & Nelson, B. W. (2014). Fire disturbance in Amazonian blackwater floodplain forests. Plant Ecology and Diversity, 7(1–2), 319–327. https://doi.org/10.1080/17550874.2012.716086

Franklin, J. F., Shugart, H. H., & Harmon, M. E. (1987). Tree Death as an Ecological Process. BioScience, 37(8), 550–556. https://doi.org/10.2307/1310665

Furch, K. (1997). Chemistry of Várzea and Igapó Soils and Nutrient Inventory of Their Floodplain Forests. In The Central Amazon Floodplain: Ecology of a pulsing system (pp. 47–67). https://doi.org/10.1007/978-3-662-03416-3_3

Fva, F. V. A. (1998). Plano de Manejo do Parque Nacional do Jaú (I. B. D. M. A. E. D. R. N. R. Ibama, Ed.).

Gloor, M., Brienen, R. J. W., Galbraith, D., Feldpausch, T. R., Schöngart, J., Guyot, J. L., … Phillips, O. L. (2013). Intensification of the Amazon hydrological cycle over the last two decades. Geophysical Research Letters, 40(9), 1729–1733. https://doi.org/10.1002/grl.50377

Gordon, A. D., & Birks, H. J. B. (1972). Numerical methods in quaternary palaeoecology I. Zonation of pollen diagrams. New Phytologist, 71(5), 961–979. https://doi.org/10.1111/j.1469-8137.1972.tb01976.x

Gower, J. C. (1971). A General Coefficient of Similarity and Some of Its Properties. Biometrics, 27(4), 857. https://doi.org/10.2307/2528823

Granato-Souza, D., Stahle, D. W., Barbosa, A. C., Feng, S., Torbenson, M. C. A., de Assis Pereira, G., … Griffin, D. (2019). Tree rings and rainfall in the equatorial Amazon. Climate Dynamics, 52(3–4), 1857–1869. https://doi.org/10.1007/s00382-018-4227-y

Green, J. W. (1963). Wood cellulose. Methods in Carbohydrate Chemistry, 9–12. Retrieved from https://ci.nii.ac.jp/naid/10029317546/

Grill, G., Lehner, B., Thieme, M., Geenen, B., Tickner, D., Antonelli, F., … Zarfl, C. (2019). Mapping the world’s free-flowing rivers. Nature, 569(7755), 215–221. https://doi.org/10.1038/s41586-019-1111-9

Hartmann, H., Adams, H. D., Hammond, W. M., Hoch, G., Landhäusser, S. M., Wiley, E., & Zaehle, S. (2018). Identifying differences in carbohydrate dynamics of seedlings and mature trees to improve carbon allocation in models for trees and forests. Environmental and Experimental Botany, 152(March), 7–18. https://doi.org/10.1016/j.envexpbot.2018.03.011

Hartmann, H., Moura, C. F., Anderegg, W. R. L., Ruehr, N. K., Salmon, Y., Allen, C. D., … O’Brien, M. (2018). Research frontiers for improving our understanding of drought-induced tree and forest mortality. New Phytologist, 218(1), 15–28. https://doi.org/10.1111/nph.15048

Hua, Q., Barbetti, M., & Rakowski, A. Z. (2013). Atmospheric radiocarbon for the period 1950– 2010. Radiocarbon, 55(4), 2059–2072. https://doi.org/10.2458/azu_js_rc.v55i2.16177

Huang, Y., Mori, S. A., & Kelly, L. M. (2015). Toward a phylogenetic-based Generic Classification of Neotropical Lecythidaceae—I. Status of Bertholletia, Corythophora, Eschweilera and Lecythis. Phytotaxa, 203(2), 85. https://doi.org/10.11646/phytotaxa.203.2.1

Idesam, I. de C. e D. S. do A. (2009). Série Técnica Planos de Gestão Reserva de Desenvolvimento Sustentável do Uatumã Volumes 1 e 2. 1 and 2, 150.

Jackson, M. B. (1990). Hormones and developmental change in plants subjected to submergence or soil waterlogging. Aquatic Botany, 38(1), 49–72. https://doi.org/10.1016/0304-3770(90)90098-6

Johnson, E. A., & Fryer, G. I. (1989). Population dynamics in lodgepole pine-Engelmann spruce forests. Ecology. https://doi.org/10.2307/1938193

Jones, P. D., & Harris, I. C. (2008). Dataset Collection Record: Climatic Research Unit (CRU): Time-series (TS) datasets of variations in climate with variations in other phenomena v3. Retrieved December 3, 2019, from University of East Anglia - NCAS British Atmospheric Data Centre website: https://catalogue.ceda.ac.uk/uuid/3f8944800cc48e1cbc29a5ee12d8542d

Juggins, S. (2017). Rioja R Package: Analysis of Quaternary Science Data. Retrieved from http://www.staff.ncl.ac.uk/stephen.juggins/

Junk, W J. (1989). Flood tolerance and tree distribution in central Amazonian floodplains. In Tropical forests: botanical dynamics, speciation and diversity. https://doi.org/http://dx.doi.org/10.1016/B978-0-12-353550-4.50012-5

Junk, Wolfgang J., Bayley, P. B., & Sparks, R. E. (1989). The flood pulse concept in river-floodplain systems. In D. P. Dodge (Ed.), Proceedings of the International Large River Symposium (Vol. 106, pp. 110–127). https://doi.org/10.1371/journal.pone.0028909

Junk, Wolfgang J., Piedade, M. T. F., Cunha, C. N. da, Wittmann, F., & Schöngart, J. (2018). Macrohabitat studies in large Brazilian floodplains to support sustainable development in the face of climate change. Ecohydrology and Hydrobiology, 18(4), 334–344. https://doi.org/10.1016/j.ecohyd.2018.11.007

Junk, Wolfgang J., Piedade, M. T. F., Schöngart, J., Cohn-Haft, M., Adeney, J. M., & Wittmann, F. (2011). A classification of major naturally-occurring amazonian lowland wetlands. Wetlands, 31(4), 623–640. https://doi.org/10.1007/s13157-011-0190-7

Junk, Wolfgang J., Wittmann, F., Schöngart, J., & Piedade, M. T. F. (2015). A classification of the major habitats of Amazonian black-water river floodplains and a comparison with their white-water counterparts. Wetlands Ecology and Management, 23(4), 677–693. https://doi.org/10.1007/s11273-015-9412-8

Kubitzki, K., & Ziburski, A. (1994). Seed Dispersal in Flood Plain Forests of Amazonia. Biotropica, 26(1), 30–43. Retrieved from https://www.jstor.org/stable/2389108

Latrubesse, E. M. (2008). Patterns of anabranching channels: The ultimate end-member adjustment of mega rivers. Geomorphology, 101(1–2), 130–145. https://doi.org/10.1016/j.geomorph.2008.05.035

Latrubesse, E. M., Arima, E. Y., Dunne, T., Park, E., Baker, V. R., D’Horta, F. M., … Stevaux, J. C. (2017). Damming the rivers of the Amazon basin. Nature, 546(7658), 363–369. https://doi.org/10.1038/nature22333

Lees, A. C., Peres, C. A., Fearnside, P. M., Schneider, M., & Zuanon, J. A. S. (2016). Hydropower and the future of Amazonian biodiversity. Biodiversity and Conservation, 25(3), 451–466. https://doi.org/10.1007/s10531-016-1072-3

Li, S., Lens, F., Espino, S., Karimi, Z., Klepsch, M., Schenk, H. J., … Jansen, S. (2016). INTERVESSEL PIT MEMBRANE THICKNESS AS A KEY DETERMINANT of EMBOLISM RESISTANCE in ANGIOSPERM XYLEM. IAWA Journal, 37(2), 152–171. https://doi.org/10.1163/22941932-20160128

Ligon, F. K., Dietrich, W. E., & Trush, W. J. (1995). Downstream Ecological Effects of Dams. BioScience, 45(3), 183–192. https://doi.org/10.2307/1312557

Lobo, G. de S., Wittmann, F., & Piedade, M. T. F. (2019). Response of black-water floodplain (igapó) forests to flood pulse regulation in a dammed Amazonian river. Forest Ecology and Management, 434, 110–118. https://doi.org/10.1016/j.foreco.2018.12.001

Lovejoy, T. E., & Nobre, C. (2018). Amazon Tipping Point. Science Advances, 4(2), eaat2340. https://doi.org/10.1126/sciadv.aat2340

Maechler, M., Rousseeuw, P., Struyf, A., Hubert, M., & Hornik, K. (2019). R Cluster Package: Cluster Analysis Basics and Extensions.

Maia, L., & Piedade, M. T. F. (2000). Phenology of Eschweilera tenuifolia (Lecythidaceae) in Flooded Forest of the Central Amazonia - Brazil. In R. Lieberei, H. K. Bianchi, V. Boehm, & C. Reisdorff (Eds.), German-Brazilian Workshop on Neotropical Ecosystems – Achievements and Prospects of Cooperative Research (p. 5). Retrieved from http://hdl.handle.net/11858/00-001M-0000-000F-DDEC-2

Malhi, Y., Roberts, J. T., Betts, R. a, Killeen, T. J., Li, W., & Nobre, C. a. (2008). Climate Change, Deforestation, and the Fate of the Amazon. Science, 319(5860), 169–172. https://doi.org/10.1126/science.1146961

Marengo, J. A., Aragão, L. E. O. C., Cox, P. M., Betts, R., Costa, D., Kaye, N., … Reis, V. (2016). Impacts of climate extremes in Brazil the development of a web platform for understanding long-term sustainability of ecosystems and human health in amazonia (pulse-Brazil). Bulletin of the American Meteorological Society, 97(8), 1341–1346. https://doi.org/10.1175/BAMS-D-14-00177.1

Marengo, J. A., Souza Jr., C. M., Thonicke, K., Burton, C., Halladay, K., Betts, R. A., … Soares, W. R. (2018). Changes in Climate and Land Use Over the Amazon Region: Current and Future Variability and Trends. Frontiers in Earth Science, 6(December). https://doi.org/10.3389/feart.2018.00228

Melack, J. M., & Hess, L. L. (2010). Remote Sensing of the Distribution and Extent of Wetlands in the Amazon Basin. In Wolfgang Johannes Junk, M. T. F. Piedade, F. Wittmann, J. Schongart, & P. Parolin (Eds.), Amazonian floodplain forests (pp. 43–59). https://doi.org/10.1007/978-90-481-8725-6_3

Mori, G. B., Schietti, J., Poorter, L., & Piedade, M. T. F. (2019). Trait divergence and habitat specialization in tropical floodplain forests trees. PLoS ONE, 14(2). https://doi.org/10.1371/journal.pone.0212232

Mori, S. A. (2001). A Família da Castanha-do-Pará : Símbolo do Rio Negro. In Florestas do Rio Negro (pp. 118–141).

Mori, S. A., & Prance, G. T. (1990). Flora Neotropica Monograph 21 (II) Lecythidaceae - part II. The zygomorphic-flowered New World genera (Couroupita, Corythophora, Bertholletia, Couratari, Eschweilera & Lecythis). In Source: Flora Neotropica Aiphanes (Palmae) (Vol. 70). https://doi.org/10.2307/2805360

Negrón-Juárez, R. I., Chambers, J. Q., Guimaraes, G., Zeng, H., Raupp, C. F. M., Marra, D. M., … Higuchi, N. (2010). Widespread Amazon forest tree mortality from a single cross-basin squall line event. Geophysical Research Letters, 37(16), 1–5. https://doi.org/10.1029/2010GL043733

Nelson, B. W. (2001). Fogo em florestas da Amazônia Central em 1997. Anais X SBSR, (June), 1675–1682.

Neves, J. R. D., Piedade, M. T. F., Resende, A. F. de, Feitosa, Y. O., & Schöngart, J. (2019). Impact of climatic and hydrological disturbances on blackwater floodplain forests in Central Amazonia. Biotropica, 51(4), 484–489. https://doi.org/10.1111/btp.12667

Nobre, C. A., Marengo, J. A., & Artaxo, P. (2009). Understanding the climate of Amazonia: Progress from LBA. Washington DC American Geophysical Union Geophysical Monograph Series, 145–147. https://doi.org/10.1029/2009GM000903

Nobre, C. A., Sampaio, G., Borma, L. S., Castilla-Rubio, J. C., Silva, J. S., & Cardoso, M. (2016). Land-use and climate change risks in the Amazon and the need of a novel sustainable development paradigm. Proceedings of the National Academy of Sciences, 113(39), 10759–10768. https://doi.org/10.1073/pnas.1605516113

Nowacki, G. J., & Abrams, M. D. (1997). Radial-growth averaging Criteria for reconstruction disturbance histories from presettlement-origin oaks. Ecological Monographs, 67(2), 225. https://doi.org/10.2307/2963514

Oliver, C. D., & Larson, B. C. (1996). Forest stand dynamics: updated edition. Forest Stand Dynamics, 520. Retrieved from https://www.cabdirect.org/cabdirect/abstract/19980604521

ONS. (2016). Atualização De Séries Históricas De Vazões – Período 1931 a 2015. Operador Nacional do Sistema – ONS. 38p.

Parolin, P. (2009). Submerged in darkness: Adaptations to prolonged submergence by woody species of the Amazonian floodplains. Annals of Botany, 103(2), 359–376. https://doi.org/10.1093/aob/mcn216

Parolin, P., De Simone, O., Haase, K., Waldhoff, D., Rottenberger, S., Kuhn, U., … Junk, W. J. (2004). Central Amazonian Floodplain Forests: Tree Adaptations in a Pulsing System. The Botanical Review, 70(3), 357–380.

Parolin, P., & Ferreira, L. (1998). Are there differences in specific wood gravities between trees in várzea and igapó (Central Amazonia). Ecotropica, 4, 25–32.

Parolin, P., & Worbes, M. (2000). Wood density of trees in black water floodplains of Rio Jaú National Park, Amazonia, Brazil. Acta Amazonica, 30(3), 441–441. https://doi.org/10.1590/1809-43922000303448

Phillips, O. L., Aragão, L. E. O. C. O. C., Lewis, S. L., Fisher, J. B., Lloyd, J., López-González, G., … Vargas, P. N. (2009). Drought Sensitivity of the Amazon Rainforest. Science, 323(5919), 1344–1347. https://doi.org/10.1126/science.1164033

Piedade, M. T. F. F., Ferreira, C. S., Wittmann, A. de O., Buckeridge, M. S., Parolin, P., Oliveira-Wittmann, A., … Parolin, P. (2010). Biochemistry of Amazonian floodplain trees. In Amazonian floodplain forests: ecophysiology, biodiversity and sustainable management (pp. 127–140). https://doi.org/10.1007/978-90-481-8725-6_6

Piedade, M. T. F., Schongart, J., Wittmann, F., Junk, W. J., & Parolin, P. (2013). Impactos ecológicos da inundação e seca na vegetação das áreas alagáveis amazônicas. In L. S. Borma & C. A. Nobre (Eds.), Secas na Amazônia: causas e conseqüências (pp. 405–457).

Prance, G. T. (1980). A terminologia dos tipos de florestas amazônicas sujeitas a inundação. Acta Amazonica, 10(3), 495–504. https://doi.org/http://dx.doi.org/10.1590/1809-43921980103499

R Core Team. (2018). R: A Language and Environment for Statistical Computing. Retrieved from https://www.r-project.org/

Ramsey, C. B. (2001). Development of the radiocarbon calibration program. Radiocarbon, 43(2), 355–363. https://doi.org/papers2://publication/uuid/5B399323-3576-4144-BF98-497069142583

Resende, Angélica F., Nelson, B. W., Flores, B. M., de Almeida, D. R., de Resende, A. F., Nelson, B. W., … de Almeida, D. R. (2014). Fire damage in seasonally flooded and upland forests of the Central Amazon. Biotropica, 46(6), 643–646. https://doi.org/10.1111/btp.12153

Resende, Angélica Faria de, Schöngart, J., Streher, A. S., Ferreira-Ferreira, J., Piedade, M. T. F., & Silva, T. S. F. (2019). Massive tree mortality from flood pulse disturbances in Amazonian floodplain forests: The collateral effects of hydropower production. Science of The Total Environment, 659, 587–598. https://doi.org/10.1016/j.scitotenv.2018.12.208

Ropelewski, C. F., & Jones, P. D. (1987). An extension of the Tahiti–Darwin southern oscillation index. Monthly Weather Review, 115(9), 2161–2165.

Rosa, S. A., Barbosa, A. C. M. C., Junk, W. J., da Cunha, C. N., Piedade, M. T. F., Scabin, A. B., … Schöngart, J. (2017). Growth models based on tree-ring data for the Neotropical tree species Calophyllum brasiliense across different Brazilian wetlands: implications for conservation and management. Trees - Structure and Function, 31(2), 729–742. https://doi.org/10.1007/s00468-016-1503-5

Sauter, M. (2013). Root responses to flooding. Current Opinion in Plant Biology, 16(3), 282–286. https://doi.org/10.1016/j.pbi.2013.03.013

Schlüter, U. B., & Furch, B. (1992). Morphological, anatomical, and physiological investigations on the tolerance to flooding by the tree Macrolobium acaciifolium, characteristic of the white- and blackwater inundation forest near Manaus, Amazonas. 12(1), 51–69. Retrieved from http://www.scopus.com/inward/record.url?eid=2-s2.0-0027088341&partnerID=40&md5=28b12d6f44e7a841bfd2cd4b299a144c

Schöngart, J., Bräuning, A., Barbosa, A. C. M. C., Lisi, C. S., & de Oliveira, J. M. (2017). Dendroecological Studies in the Neotropics: History, Status and Future Challenges. In Dendroecology (pp. 35–73). https://doi.org/10.1007/978-3-319-61669-8_3

Schöngart, J., Gribel, R., Ferreira da Fonseca-Junior, S., Haugaasen, T., Group, P. P., Cient, D. D. P., … Management, N. R. (2015). Age and Growth Patterns of Brazil Nut Trees (Bertholletia excelsa Bonpl.) in Amazonia, Brazil. Biotropica, 47(5), 550–558. https://doi.org/10.1111/btp.12243

Schöngart, J., Junk, W. J., Piedade, M. T. F., Ayres, J. M., Hüttermann, A., & Worbes, M. (2004). Teleconnection between tree growth in the Amazonian floodplains and the El Niño-Southern Oscillation effect. Global Change Biology, 10(5), 683–692. https://doi.org/10.1111/j.1529-8817.2003.00754.x

Schöngart, J., Piedade, M. T. F., Ludwigshausen, S., Horna, V., & Worbes, M. (2002). Phenology and stem-growth periodicity of tree species in Amazonian floodplain forests. Journal of Tropical Ecology, 18(4), 581–597. https://doi.org/10.1017/S0266467402002389

Schöngart, J., Piedade, M. T. F., Wittmann, F., Junk, W. J., & Worbes, M. (2005). Wood growth patterns of Macrolobium acaciifolium (Benth.) Benth. (Fabaceae) in Amazonian black-water and white-water floodplain forests. Oecologia, 145(3), 454–461. https://doi.org/10.1007/s00442-005-0147-8

Schöngart, J., Wittmann, F., Junk, W. J., & Piedade, M. T. F. (2017). Vulnerability of Amazonian floodplains to wildfires differs according to their typologies impeding generalizations. Proceedings of the National Academy of Sciences, 114(41), E8550–E8551. https://doi.org/10.1073/pnas.1713734114

Schweingruber, F. H. (1996). Principles of Dendrochronology. (July 2015).

Sevanto, S., Mcdowell, N. G., Dickman, L. T., Pangle, R., & Pockman, W. T. (2014). How do trees die? A test of the hydraulic failure and carbon starvation hypotheses. Plant, Cell and Environment, 37(1), 153–161. https://doi.org/10.1111/pce.12141

Showman, A. P., Wordsworth, R. D., Merlis, T. M., & Kaspi, Y. (2013). Atmospheric Circulation of Terrestrial Exoplanets. Radiocarbon. https://doi.org/110.2458/azu_js_rc.v51i1.3494

Sioli, H. (1984). The Amazon and its main affluents: Hydrography, morphology of the river courses, and river types. 127–165. https://doi.org/10.1007/978-94-009-6542-3_5

Speer, J. H. (2009). Fundamentals of Tree-Ring Research.

Steinhof, A., Altenburg, M., & Machts, H. (2017). Sample Preparation at the Jena 14C Laboratory. Radiocarbon, 59(3), 815–830. https://doi.org/10.1017/RDC.2017.50

Steinhof, Axel. (2013). Data Analysis at the jena 14C Laboratory. Proceedings of the 21st International Radiocarbon Conference, 55(2–3), 282–293. https://doi.org/10.2458/azu

ter Steege, H., Henkel, T. W., Helal, N., Marimon, B. S., Marimon-Junior, B. H., Huth, A., … Melgaço, K. (2019). Rarity of monodominance in hyperdiverse Amazonian forests. Scientific Reports, 9(1). https://doi.org/10.1038/s41598-019-50323-9

Timpe, K., & Kaplan, D. (2017). The changing hydrology of a dammed Amazon. Science Advances, 3(11), 1–14. https://doi.org/10.1126/sciadv.1700611

Trouet, V., & Van Oldenborgh, G. J. (2013). KNMI Climate Explorer: A Web-Based Research Tool for High-Resolution Paleoclimatology. Tree-Ring Research, 69(1), 3–13. https://doi.org/10.3959/1536-1098-69.1.3

Urli, M., Porté, A. J., Cochard, H., Guengant, Y., Burlett, R., & Delzon, S. (2013). Xylem embolism threshold for catastrophic hydraulic failure in angiosperm trees. Tree Physiology, 33(7), 672–683. https://doi.org/10.1093/treephys/tpt030

Veblen, T. T. (1986). Age and Size Structure of Subalpine Forests in the Colorado Front Range. Bulletin of the Torrey Botanical Club, 113(3), 225–240. https://doi.org/10.2307/2996361

Villalba, R., & Veblen, T. T. (1998). Influences of Large-Scale Climatic Variability on Episodic Tree Mortality in Northern Patagonia. Ecology, 79(8), 2624–2640.

Voesenek, L. A. C. J. (2003). Interactions Between Plant Hormones Regulate Submergence-induced Shoot Elongation in the Flooding-tolerant Dicot Rumex palustris. Annals of Botany, 91(2), 205–211. https://doi.org/10.1093/aob/mcf116

Williamson, G. B., Laurance, W. F., Oliveira, A. A., Delamônica, P., Gascon, C., Lovejoy, T. E., & Pohl, L. (2000). Amazonian tree mortality during the 1997 El Nino drought. Conservation Biology, 14(5), 1538–1542. https://doi.org/10.1046/j.1523-1739.2000.99298.x

Wittmann, F., & Junk, W. J. (2016). The Wetland Book. The Wetland Book, 1–20. https://doi.org/10.1007/978-94-007-6173-5

Wittmann, F., Schöngart, J., & Junk, W. J. (2010). Phytogeography, Species Diversity, Community Structure and Dynamics of Central Amazonian Floodplain Forests. https://doi.org/10.1007/978-90-481-8725-6_4

Worbes, M. (1989). Growth Rings, Increment and Age of Trees in Inundation Forests, Savannas and a Mountain Forest in the Neotropics. IAWA Journal, 10(2), 109–122. https://doi.org/10.1163/22941932-90000479

Worbes, M. (1997). The Forest Ecosystem of the Floodplains. Ecological Studies, 126(1985), 22. https://doi.org/10.1007/978-3-662-03416-3

Worbes, M., Klinge, H., Revilla, J. D. J. D., & Martius, C. (1992). On the dynamics, floristic subdivision and geographical distribution of várzea forests in Central Amazonia. Journal of Vegetation Science, 3(4), 553–564. https://doi.org/10.2307/3235812

Zeppel, M. J. B., Adams, H. D., & Anderegg, W. R. L. (2011). Mechanistic causes of tree drought mortality: recent results, unresolved questions and future research needs. New Phytologist, 192(4), 800–803. https://doi.org/10.1111/j.1469-8137.2011.03960.x

