## Supplementary Material for "Flood-pulse disturbances as a threat for long-living Amazonian trees"

*Further details on the field collections*

Almost all the wood samples were collected with the cambium present (the bark of this species is easily detached and lost because of its fibrous inner bark, locally named *envira*) or at least with sapwood to ensure the preservation of the last ring (or close to it). Only two dead trees were collected with bark attached (JNP). The majority of sampled living trees and all dead trees are from monodominant *Eschweilera*-populations in lakes, only a small number of living individuals where additionally collected from mixed forests along tributaries at the low igapó elevations (13 from JNP and 12 from USDR). Most of the sampling occurred during the terrestrial phase (September to December) of 2015, some dead trees were sampled at the USDR in 2016 and two stem discs of living trees in 2018.

From all trees, we obtained the geographic position by GPS, measured the diameter at breast height (DBH) and the flood height printed on the trunk from the last maximum water level. Based on the flood-height we estimate the elevation of the topography, using daily water-level records from nearby hydrological stations of the network from the Brazilian Water Agency (Agência Nacional de Águas – ANA). For the JNP we used the station of Manaus port (id code: 14990000; period: 1902-2018, which presented a r of 0.98 correlation with the hydrological stations Seringalzinho and Carabinani, both within the Jaú River basin, but with short periods of sampling and missing data). For the USDR the station Cachoeira Morena (id code:16100000; period: 1973-2018) in the igapó downstream of the Balbina dam was used to calculate the duration of the terrestrial period of each year. The mean flood level measured for the trees sampled at JNP was 7.8 m ± 0.7 m (6.5 m to 9 m), corresponding to a flood duration from six to ten months induced by a regular flood-pulse. At the USDR the mean flood level was similar to the population at the JNP with 7.7 ± 0.6 m (6 to 8 m).

The samples were deposited at the Dendroecological Laboratory of the National Institute for Amazon Research (INPA), where they were carefully glued on appropriate wooden supports and sanded gradually up to sandpaper with 1200 grit to enable the macroscopical analysis of the growth rings, characterized by an alternation of fiber and parenchyma bands, typical for Lecythidaceae (Worbes, 2002). The first step of this analysis comprises the delimitation of growth rings under a LEICA-MS5 microscope, where all tree rings were marked using an ultrafine-tip pencil. In a second step, ring width was measured to the nearest 0.01 mm precision using a digital measuring device (LINTAB) connected to a computer with the software Time Series Analyzes and Presentation (TSAP-Win) which produced ring-width time series simultaneously to its measurement (Rinntech, Germany). Since the formation of growth rings in the Central Amazonian floodplains occurs during the growing season initiating in the second half of the year and finishing during the first half of the following year (Schöngart *et al.*, 2002) we conventionally attribute the calendar year of latewood formation to the tree ring (for example, the growth ring formed in the period 1982/83 is associated to the calendar year 1983).

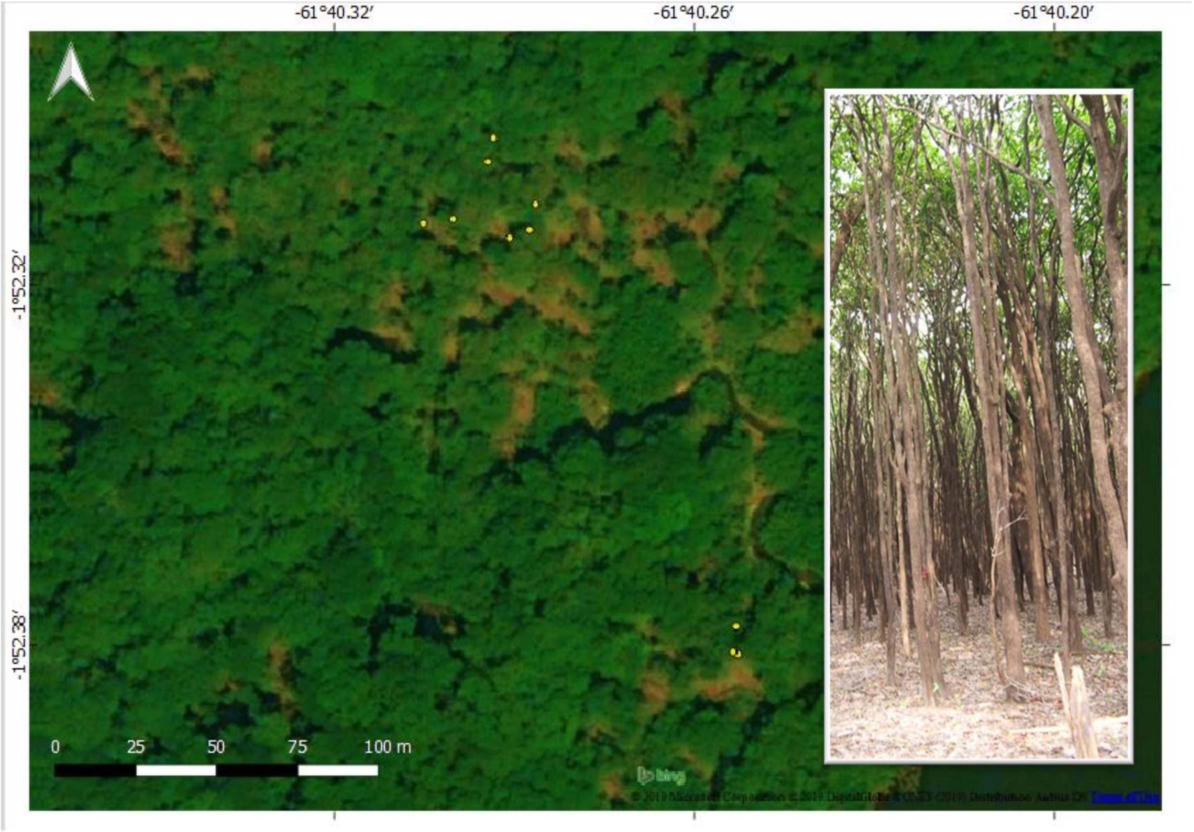


*Figure S1 - Living Eschweilera tenuifolia trees (yellow dots) at a tributary of the Jaú National Park. The crowns are closer to other tree crowns from different species.*

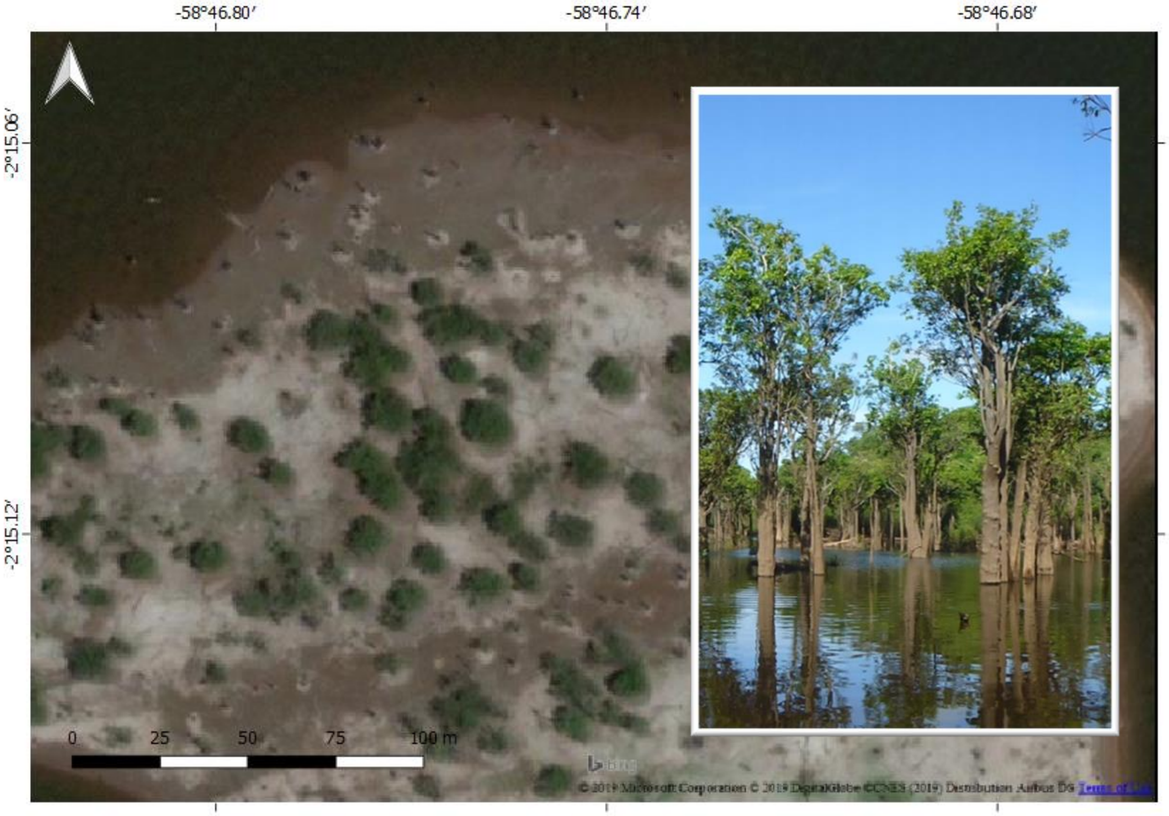


*Figure S2 – Monodominant formation of Eschweilera tenuifolia trees. All the visible crowns are living individuals of E. tenuifolia, at a lake island at the Uatumã River.*

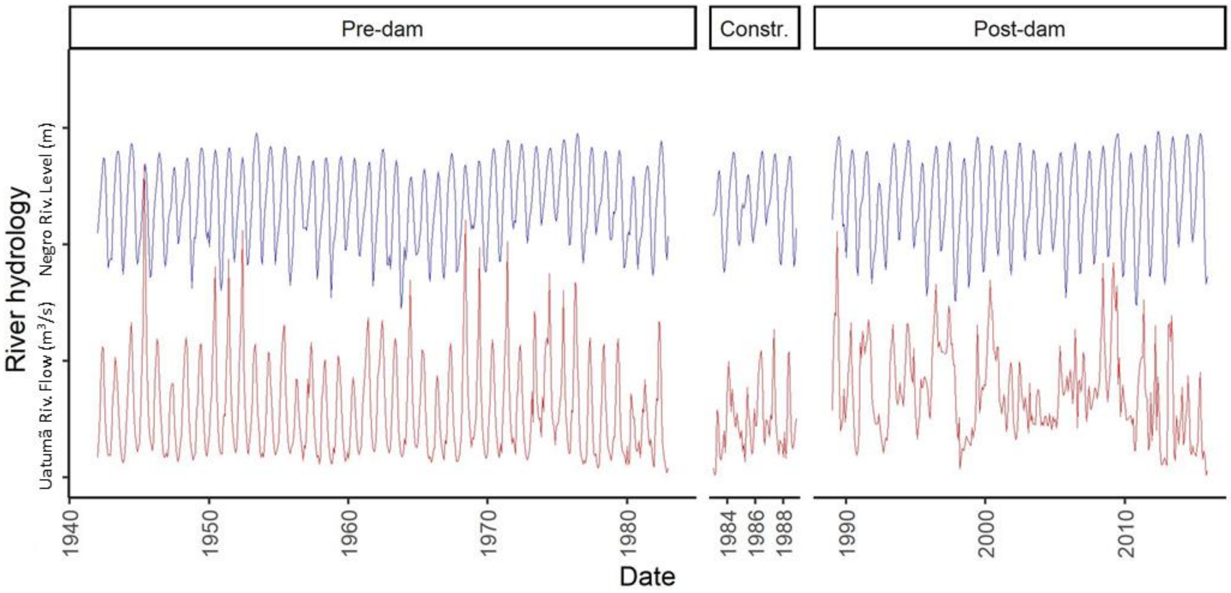


*Figure S3 – The figure illustrates the hydrology of both studied basins, the Negro river basin (in blue), representing the Jaú River and at the pristine study area and the Uatumã river basin (in red) representing the disturbed site downstream of the Balbina dam, both for the periods pre- and post-dam considering also the construction period (1983-1989). The hydrology is represented by the river level data from Negro River and discharge data for the Uatumã River. The difference in the data is due to the availability of data during the given period (both data are from the Brazilian National Agency of Waters ANA, http://www.snirh.gov.br/hidroweb/mapa)*

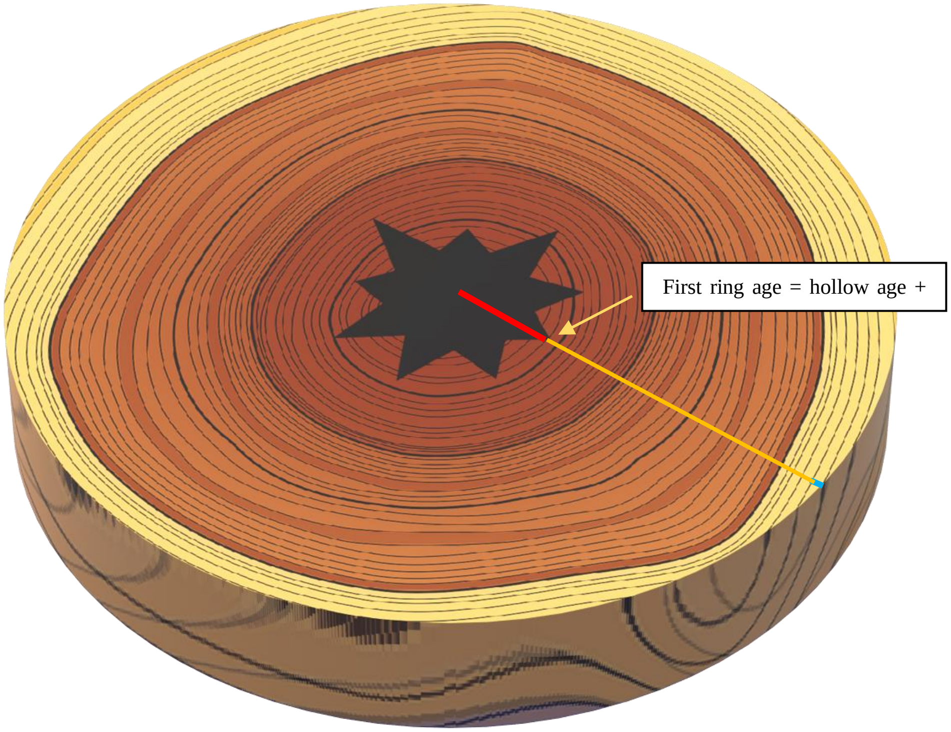


*Figure S4 - Stem disc representing a hollow tree of Eschweilera tenuifolia. The line in blue indicates the bark thickness, the line in yellow the core radius (containing heartwood and sapwood) and the red line the radius in the hollow section. To estimate the age of the hollow tree we used the age-DBH modeling (red line), adding the age of the analyzed intact wood section along two to four radii (yellow line).*

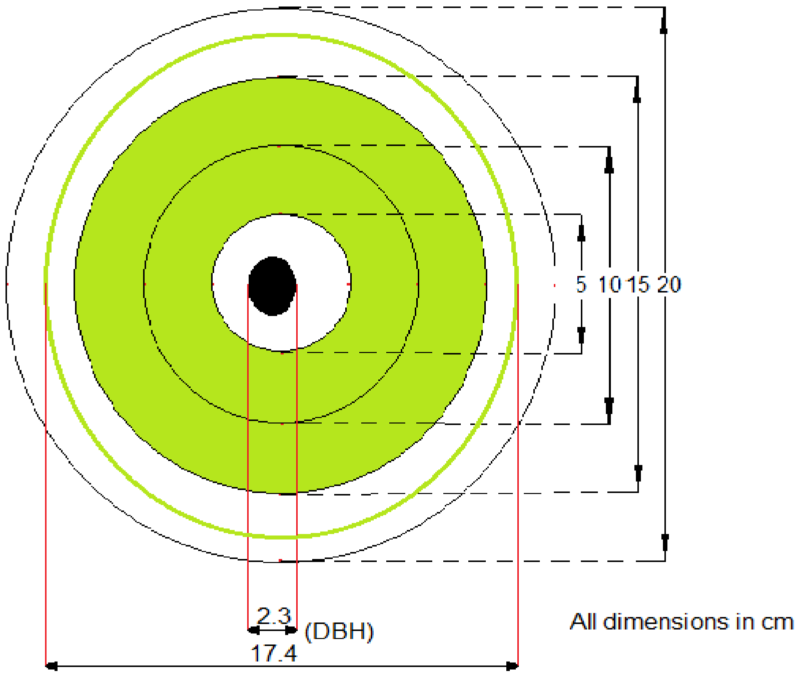


*Figure S5 - Example of how the mean passage time through 5-cm diameter classes was calculated showing an imaginary tree with 17.4 cm diameter at breast height (DBH) and a hollow of 2.3 cm (black). Only complete sections of 5-cm classes were accounted for.*

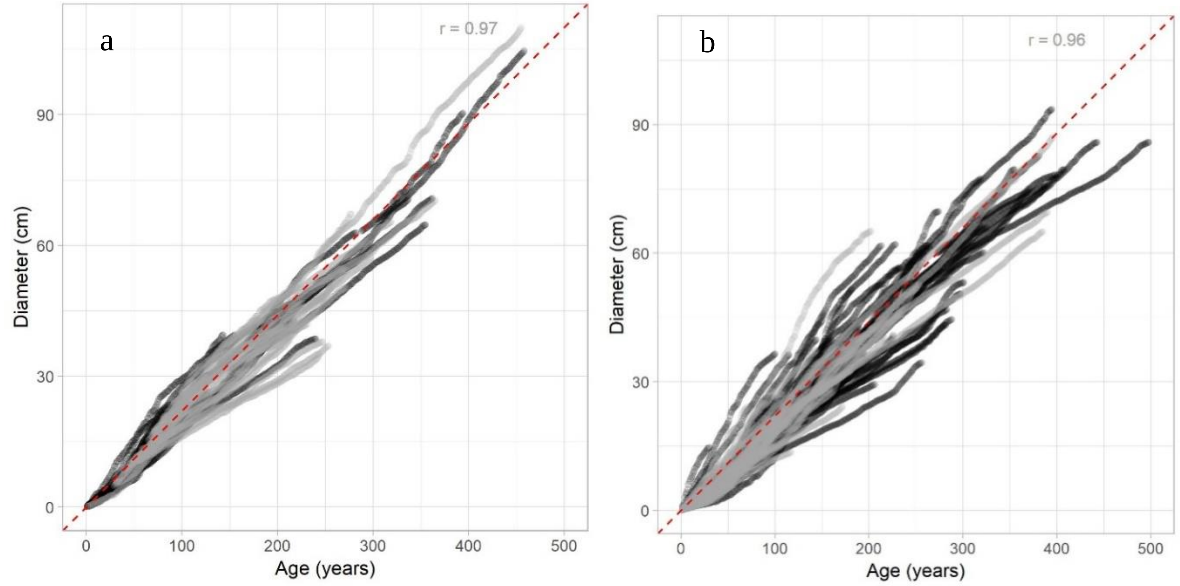


*Figure S6 – Age-diameter relationship of living (grey) and dead (black) trees of Eschweilera tenuifolia from (A) the Jaú National Park (JNP) and (B) Uatumã Sustainable Development Reserve (USDR). The red dashed line is an imaginary reference representing the same inclination for both to allow comparison.*

*Table S1 – Suppression and release events per area (Jaú National Park-JNP; Uatumã Sustainable Development Reserve-USDR), considering different macrohabitats as well as living and dead trees (no dead trees were collected in tributaries).*

| ***Macrohabitat*** | | ***Lake*** | | ***Tributary*** | |
|---|---|---|---|---|---|
| ***Area*** | Event | Living | Dead | Living | Dead |
| ***USDR*** | Release | 2 (4.0%) | 3 (10.3%) | 1 (8.3%) | -- |
| | Suppression | 2 (4.0%) | 0 | 4 (33.3%) | -- |
| | Both | 1 (2.0%) | 2 (6.9%) | 2 (16.6%) | -- |
| ***JNP*** | Release | 0 | 1 (4.8%) | 1 (7.7%) | -- |
| | Suppression | 0 | 1 (4.8%) | 2 (15.4%) | -- |
| | Both | 0 | 0 | 1 (7.7%) | -- |

*Table S2 – The tested differences between key time periods of equal length (33 years) and study sites (Jaú National Park-JNP; Uatumã Sustainable Development Reserve-USDR). Values in bold are significantly different considering Wilcox Mann-Whitney's test for at probability of 99%, while * indicates difference at 95% probability (P1: 1919-1950; P2: 1951-1982; P3: 1983-2015).*

| *Group 1* | *Group 2* | *Wilcox* | *p.value* |
|---|---|---:|---:|
| JNP_P1 | USDR_P1 | 529 | *0.849* |
| JNP_P2 | USDR_P2 | 556 | *0.889* |
| **JNP_P3** | **USDR_P3** | **296** | ***0.001*** |
| JNP_P1 | JNP_P2 | 517 | *0.731* |
| JNP_P1 | JNP_P3 | 348 | *0.01** |
| JNP_P2 | JNP_P3 | 384 | *0.04** |
| USDR_P1 | USDR_P2 | 513 | *0.693* |
| **USDR_P1** | **USDR_P3** | **220** | ***0.000*** |
| **USDR_P2** | **USDR_P3** | **256** | ***0.000*** |

*Table S3 – Radiocarbon dating results (Calendar dating using Oxcal v3.4 of SH 3 zone).) for death Eschweilera tenuifolia trees from the Jaú National Park (JNP) and Uatumã Sustainable Development reserve (USDR) considering those tree within the modern carbon period.*

| *Study Area* | *Sample code* | *Postulated period* | *14C cal. Date* | *Prob. (%)* |
|---|---|---|---|---|
| *USDR* | USDR125 | 1985 | 1982-1985 | 91.5 |
| *USDR* | USDR135 | 1989 | 1987-1990 | 63.2 |
| *USDR* | USDR136 | 1995 | 1990-1994 | 81.8 |
| *USDR* | USDR137 | 1995 | 1994-1996 | 81.7 |
| *USDR* | USDR138 | 1984 | 1980-1982 | 90.6 |
| *USDR* | USDR139 | 1999 | 1998-2000 | 92.0 |
| *USDR* | USDR140 | 1994 | 1993-1995 | 85.3 |
| *USDR* | USDR141 | 2004 | 2003-2005 | 87.6 |
| *USDR* | USDR144 | 1996 | 1997-2000 | 92.5 |
| *USDR* | USDR207 | 1986 | 1984-1986 | 74.3 |
| *USDR* | USDR212 | 1994 | 1993-1995 | 83.6 |
| *USDR* | USDR213 | 1997 | 1993-1995 | 83.4 |
| *USDR* | USDR214 | 1994 | 1993-1996 | 83.8 |
| *USDR* | USDR216 | 1984 | 1984-1985 | 61.4 |
| *USDR* | USDR217 | 1988 | 1987-1990 | 73.3 |
| *USDR* | USDR218 | 2007 | 1990-1994 | 83.3 |
| *USDR* | USDR219 | 1990 | 1989-1992 | 69.2 |
| *USDR* | USDR221 | 1995 | 1994-1996 | 83.8 |
| *USDR* | USDR238 | 1996 | 1995-1998 | 88.0 |
| *USDR* | USDR239 | 2001 | 1995-1999 | 90.8 |
| *USDR* | USDR240 | 1984 | 1982-1984 | 95.4 |
| *USDR* | USDR241 | 1993 | 1993-1996 | 83.2 |
| *USDR* | USDR242 | 1985 | 1985-1987 | 75.2 |
| *USDR* | USDR243 | 1997 | 1996-1999 | 87.9 |
| *USDR* | USDR244 | 1991 | 1989-1995 | 84.8 |
| *USDR* | USDR246 | 1975 | 1974-1976 | 91.9 |
| *JNP* | JNP082 | 2012 | 2009-2014 | 92.2 |
| *JNP* | JNP088 | 2011 | 2007-2013 | 88.8 |
| *JNP* | JNP092 | 2006 | 2010-2014 | 60.9 |
| *JNP* | JNP101 | 1974 | 1975-1976 | 77.3 |
| *JNP* | JNP108 | 2008 | 1998-2001 | 88.8 |
| *JNP* | JNP109 | 1999 | 1997-1999 | 90.5 |
| *JNP* | JNP115 | 1972 | 1972-1974 | 95.4 |
| *JNP* | JNP116 | 2010 | 2008-2011 | 90.3 |
| *JNP* | JNP121 | 1972 | 1977-1980 | 95.4 |
